# The dynamics of thermal stress events determine whether thermotolerant endosymbionts improve or hamper survival of coral reefs

**DOI:** 10.1101/2025.09.18.677250

**Authors:** A. Bonforti, F. Ohlsson, E. Libby, N. A. Kamenos

**Affiliations:** Department of Ecology, Environment and Geoscience, Umeå University, Umeå, Sweden; Integrated Science Lab, Umeå University, Umeå, Sweden; Department of Mathematics and Mathematical Statistics, Umeå University, Umeå, Sweden; Umeå Marine Sciences Centre, Umeå University, Norrbyn, Sweden

## Abstract

Coral reefs are increasingly threatened by marine heatwaves which can cause devastating coral bleaching events. Despite this, corals can sometimes survive often altering their symbiont composition toward more thermally-tolerant taxa in response to thermal stress. The long-term efficacy of such a strategy remains poorly understood, in particular, the function of bleaching as an acclimation agent. Here, we use a model of coral-symbiont dynamics, which includes novel and empirically intractable aspects of symbiont thermal tolerance, to investigate how changes in coral endosymbiont composition influence reef survival under differing patterns of heat stress. We find that thermal stress-induced bleaching can act as an acclimation mechanism, but its effectiveness is modulated by the interplay between thermal stress intensity and frequency. When thermal stress remains moderate in frequency and intensity, acclimation can occur as longer intervals between stress events promote post-bleaching coral recovery via a beneficial sequence of hosting thermally-tolerant then thermally-sensitive symbionts, interspersing thermal resilience with periods of faster coral growth. In contrast, when thermal stress intensity increases rapidly and occurs at higher frequency, sustained dominance of thermally-tolerant symbionts is favoured, providing a temporary survival strategy that delays reef collapse but with the trade-off of lower coral growth. While in this latter scenario, acclimation may be less effective due to disrupted reef population structure, it does provide a mechanism for corals to “buy time”, during which other adaptive processes may occur enabling longer-term coral survival. However, under a real-world low emission IPCC future scenario, even this flexible survival strategy may still lead to an immediate and sustained fall in coral cover. By capturing both, 1) thermal frequency and intensity dynamics, along with, 2) symbiont thermotolerance preferences and linked growth trade-offs we provide a coral survival model which can be used as a tool to inform strategic coral reef management frameworks promoting reef resilience.

## 1 Introduction

Coral bleaching occurs when the symbiotic relationship between corals and their photosynthetic algal zooxanthellae breaks down. This breakdown is often due to environmental stressors including elevated sea temperatures and high irradiance [e.g. 1, 2]. Indeed, mass bleaching events observed over the past few decades have been largely attributed to increased sea surface temperatures and marine heat waves [3] This breakdown in symbiosis leads to the expulsion of a large proportion of the zooxanthellae community, causing the coral to appear white (bleached), leading to partial or complete mortality of the coral colony [e.g. 4]. The loss of symbionts severely reduces the coral’s energy intake as zooxanthellae photosynthesis accounts for up to 90% of the holobiont’s energy needs [5]. This energy deficit impairs vital processes including growth, reproduction, and immune responses, increasing susceptibility to disease and mortality [3]. While bleaching-induced symbiont loss is often considered detrimental to coral survival, including causing reduced recovery rates more widely [6], recent evidence suggest that corals can acquire thermal resilience following stress events, which can last for at least a few years [7, 8]. In some cases, bleaching may even serve as a driver of adaptive and acclimatory processes, allowing corals to survive in changing environments [8-11]. However, whether bleaching ultimately leads to mortality or long-term resilience remains equivocal.

To determine when, or if, bleaching can lead to increased resilience, it is useful to identify a mechanism by which bleaching may lead to acclimatization. One possible mechanism of acclimatization involves altering the composition of endosymbiotic algae to favour species with greater thermal tolerance. By expelling their existing symbiont community and acquiring new symbionts from the environment, or repopulating from those that remain, corals may rapidly and efficiently swap symbionts so as to enhance their resilience to thermal stress [10, 12]. Similar means of acclimatization have been observed within the coral’s lifespan through phenotypic plasticity [13] either through symbiont reshuffling — i.e., the gradual reorganization of symbiont communities within the coral host — or symbiont switching — i.e., the acquisition of entirely new symbiont species from the environment [14]. Acclimatization can also occur across generations, through non-genetic transfer, such as vertical transmission of symbionts from parent to offspring [e.g. 15] though the benefits of this process likely take longer to manifest. Changing endosymbiotic composition is not necessarily permanent, as corals may revert to their original, stress-sensitive symbionts over time assuming no further stress events occur [16]. Thus, the overall biogeography of symbiont populations found with coral hosts is thought to be environmentally (thermally)-, rather than host-driven [17]. Shifting between symbionts may come at a cost, potentially reducing coral growth or reproductive success [e.g. 18]. Over longer timescales, these thermally-induced costs could have significant consequences on reef-scale dynamics and colony resilience [e.g. 19]. While the role of symbionts conferring thermal tolerance at the colony level is well established, the interplay between symbiont dominance and recurrent heat stress and how it translates into broader ecosystem outcomes is less well understood. It is clear that the frequency and intensity of heatwaves, along with the ability of corals to adjust their symbiont composition in response to these stressors, will shape reef trajectories under future climate conditions [7, 8].Capturing the complex interplay of these factors across different timescales is a task well suited for computational modeling. Yet, despite its potential, few modeling efforts explicitly account for how these mechanisms combine to drive either reef decline or long-term persistence.

Here, we use a mechanistic reef model, extending previous models by Mumby et al. [20] and Ortiz, et al. [21] by including reversible switching between thermally-sensitive and thermally-tolerant symbionts. We investigate how the ability of corals to adjust their symbiont composition — favoring either thermally-sensitive or thermally-tolerant symbionts — affects the fate of the reef under different regimes of heat stress. In our model we characterize recurring thermal stress based on its frequency and relative intensity. We find that the resilience of coral communities and the underlying reef dynamics are influenced by two factors: 1) whether the stress allows enough time for corals to revert to thermally-sensitive symbiont communities, and 2), whether the intensity of stress remains constant or increases over time. The interaction between these factors and bleaching-initiated symbiont swapping determines whether coral resilience improves or declines; indeed under a future IPCC scenario our model suggests an immediate decline. Notably, the dominance of thermotolerant symbionts can drive the system toward either outcome. Ultimately, our modeling framework identifies key characteristics of recurring thermal stress that predict when and under what conditions bleaching-induced symbiont swapping may enhance reef survival.

## 2. Summary Methods

### 2.1 Model overview

Elegant coral metacommunity modelling frameworks exist (e.g. *C∼scape* [22]) which model coral dynamics, although at present do not incorporate the direct role of symbionts in driving those dynamics. Also, dynamic energy budget models of symbiont competition on coral colony host survival exist (e.g. Cunning et al. [23] and Brown et al. [24]), but generally cannot incorporate reef scale survival trajectories to projected warming. To bridge these gaps, we incorporate reversible symbiont-endowed host thermal tolerance, thermal environmental trajectories and reef wide coral dynamics by building upon an established coral reef model developed by Mumby et al. [20] and expanded by Ortiz et al. [21]. The Mumby-Ortiz model was designed for mid-depth forereefs, and we adapted it to explore how reversible switching between thermally-sensitive and thermally-tolerant symbionts affects reef dynamics under repeated thermal stress. The model represents a spatially explicit square lattice of discrete patches, each containing live coral, macroalgae, cropped algae or unoccupied substrate. Processes including coral growth, mortality, grazing, and benthic competition (coral-coral, macroalgae-coral) are simulated with semi-annual (six-month) resolution. Grazing intensity in our model is fixed at 80% to keep macroalgae populations in check, thereby simulating a starting healthy reef. There are no disturbances besides thermal stress (e.g., no hurricanes or fishing), in order to focus on how the frequency and intensity of thermal stress events interact with symbiont exchange to shape coral trajectories. The general model mechanisms are outlined below with full details in the SM.

### 2.2 Symbiont types, switching, and reversal

When corals dominated by sensitive symbionts experience a thermal stress event, a fraction of the surviving colonies will probabilistically switch to the tolerant symbiont. The probability of switching depends on the intensity of the thermal stress event. If a thermal stress event occurs when the colony is dominated by tolerant symbionts, it simply prolongs their dominance. If no further thermal stress events occur during a number of consecutive six-month periods in our model, all tolerant symbiont-dominated corals immediately and deterministically revert to sensitive symbionts. This assumption is motivated by evidence that corals can revert to faster-growing symbionts once stress subsides [16, 25]. We use a deterministic reversion process in order to explore how corals exploit growth advantages in stress-free intervals. Moreover, deterministic reversal also allows us to isolate the differential response of the reef to thermal stress frequencies that are higher or lower than the characteristic symbiont reversal time. We consider the effects of our reversal assumption by exploring alternative types of reversal, e.g. probabilistic or gradual reversal, in additional simulations presented in the SM. We note that the switching and reversal process in our model captures either ‘shuffling’ (competitive rearrangement of already existing internal symbiont populations) or true ‘switching’ (uptake of new symbionts from the environment), similar to approaches of Ortiz et al. [21]. Here, we use the term ‘switching’ to describe these processes in our model, and generally assume that shuffling is the acting mechanism.

### 2.3 Growth, mortality, and reproduction dynamics

Corals hosting tolerant symbionts grow more slowly than sensitive symbiont-dominated corals but experience reduced mortality when exposed to thermal stress [18, 26] (see Table 1). Coral mortality also depends on colony size, consistent with Mumby et al. [20] and Ortiz et al. [21]. In our model corals reproduce only via production of larvae. Adult coral colonies create larvae proportional to both their size and fecundity. At the moment of release, larvae harbour the same symbiont composition and reflect the same reversal timing as the mother coral. Fecundity can be influenced by environmental factors, particularly the presence of macroalgae in the coral’s immediate surroundings. The total larval production is then calculated by multiplying the fecundity by the colony’s area. Notably, our model allows for larval saturation, i.e., situations in which more larvae are produced than there is available space for recruitment (see SM for further details and parameter values).

**Table 1:** Summary of parametric differences between thermally-sensitive and thermally-tolerant symbionts (SeS and ToS, respectively). *Mortality*: probability of mortality for corals dominated by each symbiont type during thermal stress events. *Switching/Reversal*: probability of switching from sensitive to tolerant symbionts and vice versa (reversal) after thermal stress events (see Section SM3 for additional reversal dynamics). *Growth*: growth rates for corals dominated by each symbiont type. Note that *P*_*M*(*SeS*)_, *P*_*M*(*ToS*)_, and *P*_*S*(*SeS→ToS*)_ depend on thermal stress intensity, while *P*_*R*(*ToS→SeS*)_, *Gr*_(*SeS*)_, and *Gr*_(*ToS*)_ are fixed. In our model, corals are never fully dominated by a single symbiont, as a cryptic minority always persists, with maximum occupancy for the dominant symbiont set at 90%. Actual parameter values reflect current community composition. For detailed parameter values across stress intensities, see Table SM3 and Fig. SM1.

| Mortality | Value |
| --- | --- |
| $P_M(SeS)$ | $0.9 \times \text{event intensity}$ |
| $P_M(ToS)$ | $0.1 \times \text{event intensity}$ |
| Switching/Reversal | Value |
| $P_S(SeS \rightarrow ToS)$ | $0.8 \times \text{event intensity}$ |
| $P_R(ToS \rightarrow SeS)$ | 1 (after 4 years w/o stress) |
| Growth | Value |
| $Gr(SeS)$ | 0.9 |
| $Gr(ToS)$ | 0.1 |

A key difference compared to reference models is that, while in those models a maximum of three corals can coexist in the same cell — thereby strongly limiting recruitment opportunities—in our model, recruits can occupy all available space within a cell (e.g. by successful competition; see Table SM4 for full details of differences).

### 2.4 Thermal stress scenarios

Thermal stress is introduced in our model at regular intervals of either three or five years following a sequence of historical heat stress data for the Great Barrier Reef up to the year 2020, [e.g. 8]. The stress is characterized by an intensity value that determines symbiont-dependent mortality and symbiont switching probability (see Table SM3 and Fig. SM1). In our model we examine two contrasting regimes: *1)* an increasing-intensity scenario in which the intensity of the thermal stress events increases linearly over time after the historical series (see Figure 1), and *2)* a fixed-intensity scenario in which all thermal stress events – including those in the historical sequence – are maintained at a fixed intensity equal to that of the normalized median intensity of the 2016 thermal stress event (see section SM1.10.1). Under the increasing-intensity scenario, successive bleaching events grow more severe with time. Under the fixed-intensity scenario, all bleaching events share the same severity, approximating a scenario where corals experience repeated stress as uniform, either due to some acclimating mechanism or moderate ongoing climate change. We summarize the symbiont dynamics in the two thermal stress scenarios in Figure 2.

**Figure 1:**
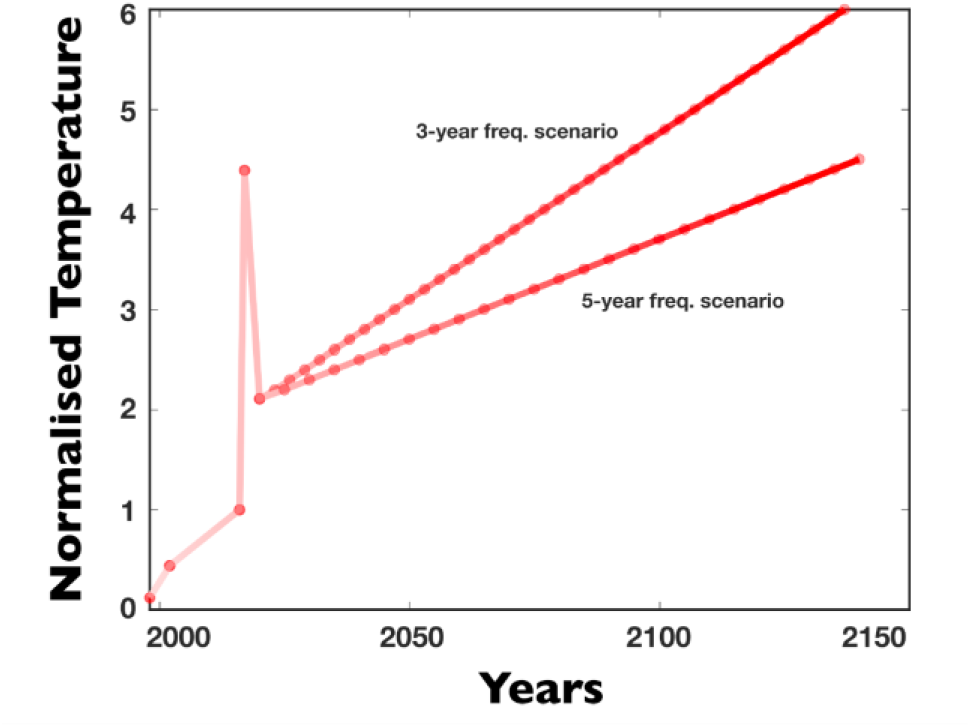
Thermal stress sequences for the increasing-intensity scenario. Thermal stress events in our model begin with a historical sequence (1998–2020) (based on the Great Barrier Reef (Low Isles), using occurrence and intensity values from Kamenos and Hennige [8]. Intensities are normalized to the 2016 event (set to 1), with all other events rescaled accordingly (see Methods and SM 1.10.1). After 2020, event intensity increases linearly. The 3-year and 5-year cycle scenarios differ in frequency and, due to fixed per-event temperature increases, also in the maximum attained temperature. For comparison, simulations excluding the historical sequence are shown in SM Section 2.

**Figure 2:**
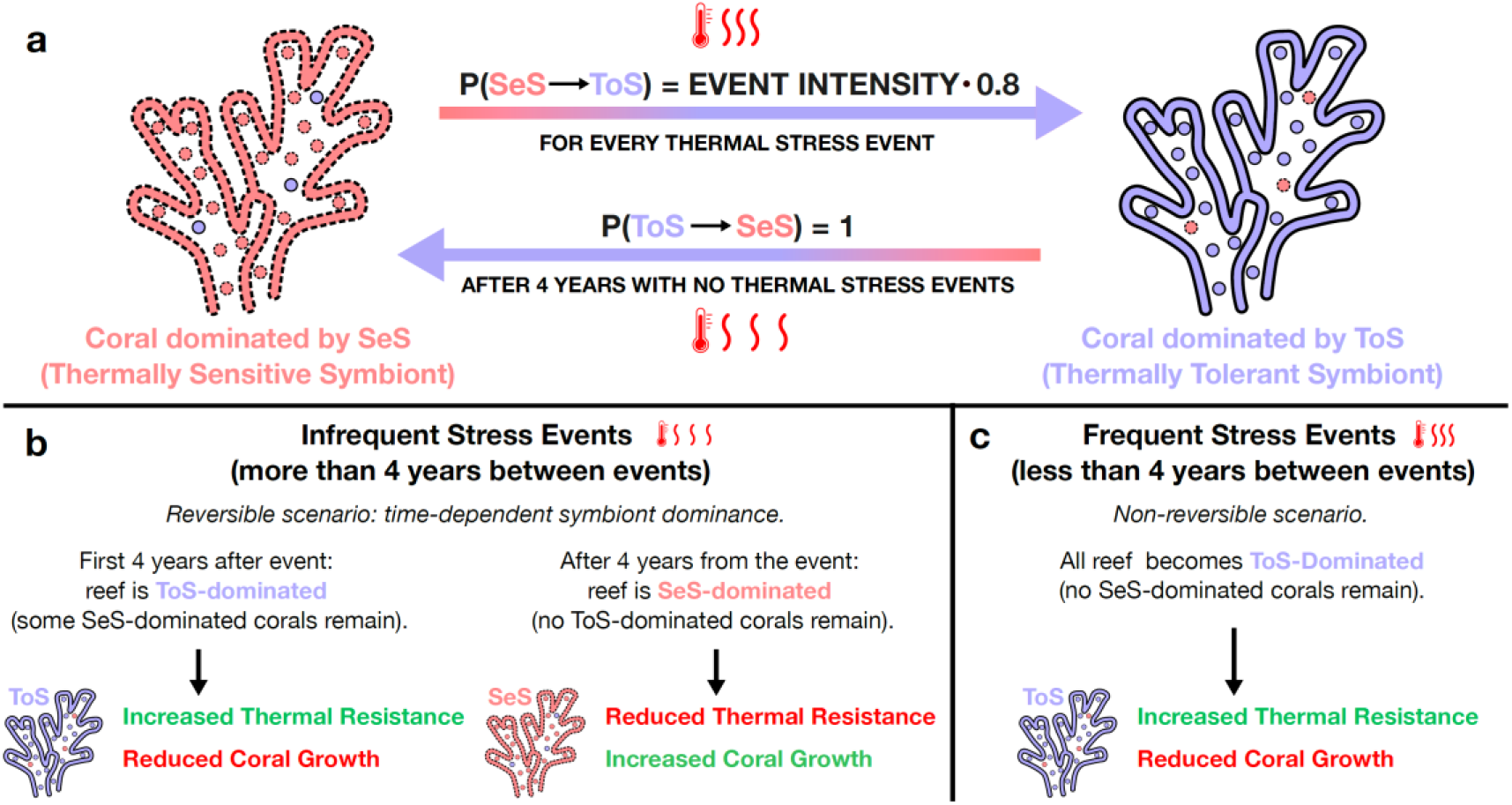
Symbiont switching and reversal dynamics. (a) The probability of switching from thermally-sensitive (SeS) to thermally-tolerant (ToS) symbionts is calculated as event intensity 0.8. For fixed thermal stress (intensity = 1), *P*_(*SeS→ToS*)_ = 0.8, while in the increasing-intensity scenario it saturates at 1 (see Table SM3 and Fig. SM1). In the main text results, reversal from ToS to SeS always occurs deterministically after 4 years without stress (see SM3.1 for alternative reversal dynamics). (b) When thermal stress events are more than 4 years apart, corals have sufficient time to revert to SeS dominance before the next event (reversible scenario). As reversal is deterministic in the main text, all corals revert fully to SeS dominance. (c) When thermal stress events occur less than 4 years apart, corals remain ToS-dominated as reversal cannot complete before the next event (non-reversible scenario). Over time, all remaining SeS-dominated corals gradually switch to ToS dominance, leading to complete ToS dominance across the reef.

We also conduct a simulation with real-world relevance using IPCC projections (‘low-emissions’ scenario SSP-126) to 2035 imposing a future projection sequence starting from 2025. We used Low Isles on the Great Barrier reef as a model location as it has the longest observational record of bleaching — starting in 1929 — and thus provides an ecosystem where we can consider the longest record of bleaching and recovery [8, 27]. To incorporate these data into our model, we calculated Degree Heating Months (DHM) stress values with a methodology similar to [28, 29] (see SM1.10.4).

### 2.5 Simulation summary

At simulation initialization, all corals are assumed to host thermally-sensitive symbionts — a choice reflecting the notion that no major bleaching events occurred in the immediate past. Because of the initial complete dominance of thermally-sensitive symbionts an extremely severe stress event could cause immediate collapse of the reef, highlighting the sensitivity to initial conditions. We address this by priming the reef response using the sequence of historical stress events to ensure corals persist before the main simulations begin [3, 8]. Similarly, in the simulations with no initial historical sequence shown in SM2, we prime the reef using an initial sequence of sublethal thermal stress events. Six-month timesteps were chosen for computational simplicity and for consistency with Mumby [20], Mumby [30] and Ortiz [21], consolidating shorter-scale processes into semi-annual updates while emphasizing multi-decade reef development. Each run spans at least 150 years (300 timesteps), capturing multiple thermal stress episodes. Coral cover, size-structure, and symbiont composition are tracked over time. We repeat coral reef simulations with different random seeds to account for the stochastic nature of processes such as switching and mortality.

While our model is based on an established coral reef model [20, 21, 30], there are some differences in modeling assumptions and parameters.

- We introduce symbiont reversal (from tolerant to sensitive) which was not implemented previously due to difficulty in parameterizing [21].
- We include the rate of switching to tolerant symbionts as well as the effects of symbiont composition on growth and mortality.
- We do not include the effects of fishing, hurricanes, and diseases.
- We consider a single generic massive coral morphotype in order to simplify our analyses.

These changes were made in order to focus on shifts in dynamics resulting from the interactions of thermal stress events and symbiont populations (see Table SM3). Complete parametric and implementation comparison between the current model and reference models is outlined in Table SM4.

## 3 Results

### 3.1 Coral cover in the absence of future thermal stress

To establish baselines for comparisons, we conduct preliminary simulations in which there are no future instances of heat-induced thermal stress (Fig. 3). We conducted two simulations: 1) we consider a scenario where there is no history of thermal stress (Fig. 3a), and 2) we include past historical thermal stress events from 1998 until 2020, but no stress events afterwards (Fig. 3b). In both scenarios, coral cover reaches the maximum allowed occupation, which indicates that in absence of thermal stress our simulated coral reefs can achieve and maintain optimal health— even after some initial, sub-lethal, thermal stress. In the scenario without thermal stress, coral cover reaches the maximum allowed occupation, which due to high algal grazing levels in our model corresponds to full reef area coverage (excluding sand patches). In the scenario without thermal stress after 2020, the reef first experiences three sub-lethal thermal stress events for both sensitive and tolerant symbionts in 1998, 2002, and 2016 (normalized intensities: 0.12, 0.44, and 1; see Table SM3). Two additional events occur in 2017 and 2020 (normalized intensities: 4.39 and 2.11), which are lethal for thermally sensitive symbiont dominated corals but sub-lethal for thermally tolerant symbiont dominated corals. Due to probabilistic thermally sensitive to tolerant symbiont switching and the 4-year deterministic reversal in this simulation, a portion of corals remain thermally tolerant symboint-dominated during the 2017 and 2020 events

**Figure 3:**
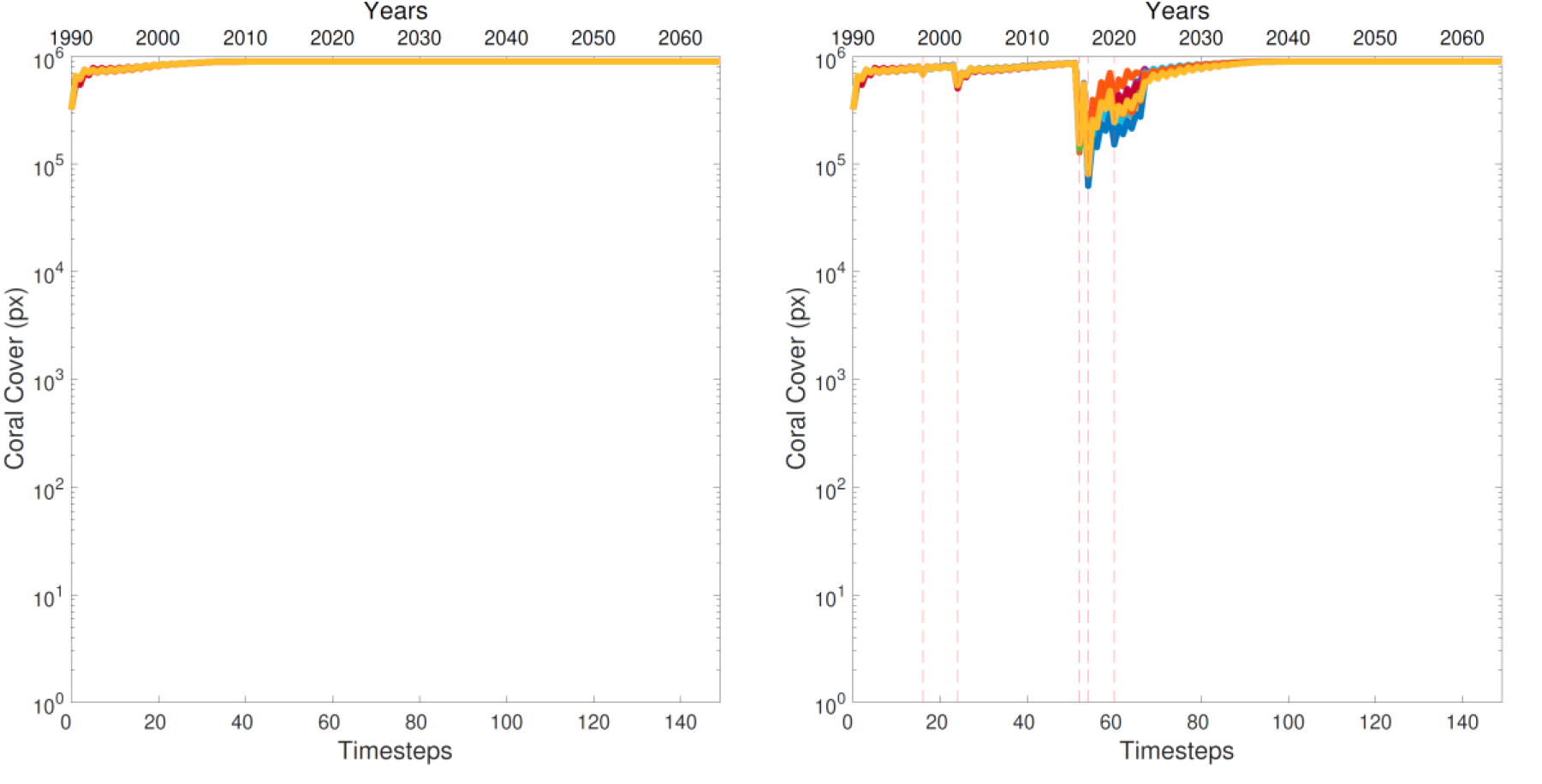
Projected coral cover (as pixels - px) over 150 simulated years under no thermal stress events (left panel), and no thermal stress events after 2020 (right panel). Ten independent runs are shown for each scenario.

### 3.2 Thermal scenario 1: Increasing future thermal stress intensity

We consider a scenario in which thermal stress events increase in intensity after the initial historical sequence of events using two model experiments.

Firstly, we explore the scenario when the thermal stress events happen at a high frequency so that there is not enough time for corals to revert to thermally sensitive symbionts (see the 3 year interval in Fig. 1). In this case when a thermal stress event occurs, the majority of sensitive symbiont-dominated corals acquire thermally tolerant symbiont-dominance (*P*_*S*(*SeS→ToS*)_ = 0.8*×*event intensity, see Table 1). As there is not enough time between stress events to allow symbiont reversal back to sensitive symbiont dominance, corals become locked into thermally tolerant symbiont-dominated states (see Fig. 4a). Corals that failed to switch to thermally tolerant symbionts are susceptible to the next stress event and experience a greater risk of death. Despite the thermally tolerant symbiont-dominated reef, our simulations show a gradual decline in coral cover that results in reef extinction after 20-40 years from the onset of high frequency thermal stress events (Fig. 4a). We note that when the corals die in these simulations the intensity of stress events is not high enough to kill the corals dominated by thermally tolerant symbionts. Instead, corals die because of a trade-off within the agent-based model between thermal protection and growth rate [e.g. 31]. While thermally tolerant symbiont-dominated corals have enhanced thermal resistance, they are not able to sustain larval growth towards adulthood and eventually die due to thermal stress. We explore this restructuring later in the Results (Fig. 6a).

**Figure 4:**
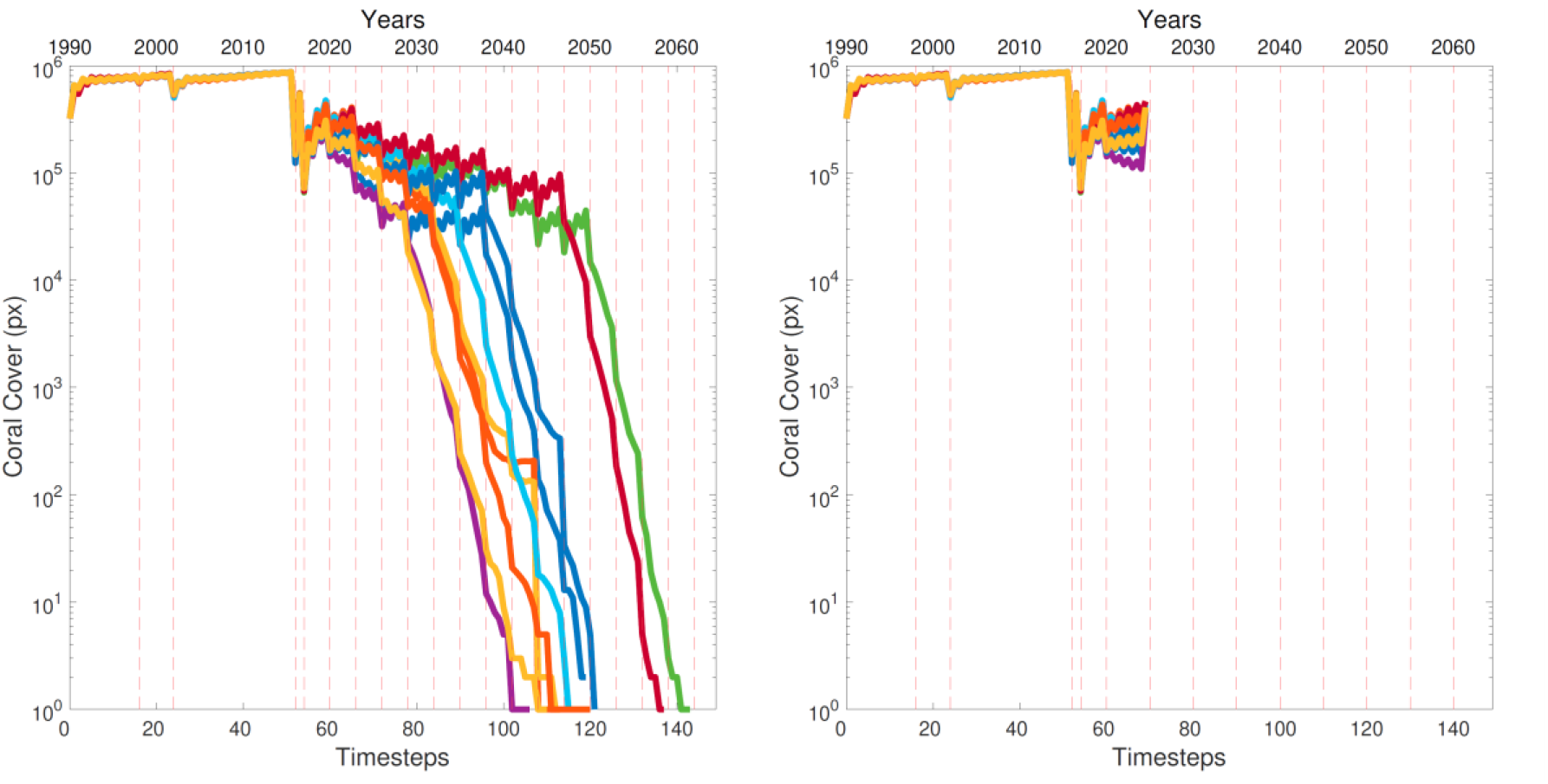
Projected coral cover (as pixels - px) over 75 simulated years (150 timesteps) under thermal stress events with increasing intensity every 3 years (a - left panel) or 5 years (b - right panel), with deterministic 4-year reversal. In the 3-year cycle (left), thermal stress events occur before thermally tolerant symbionts can revert to sensitive symbionts, so that the reef is thermally tolerant symbiont-dominated when new stress events occur. Ten independent runs are shown for each scenario.

We then explore the scenario when thermal events happen at a lower frequency so that corals can revert from thermally tolerant to sensitive symbionts (see the 5-year interval in Fig. 1). The resulting simulations show that the reef undergoes a more rapid decline than in the previous case of high frequency events (see Fig. 4b). All reefs go extinct before 35 simulated years (70 timesteps) compared to 50-70 simulated years (100-140 timesteps) in the case of high frequency events, where the total number of stress events is higher. There are two factors that cause reefs to go extinct faster when exposed to low frequency events. First, corals have sufficient time to revert to sensitive-dominated communities, which are more susceptible to death in response to the next thermal stress event. Second, because corals switch (with some probability) to the tolerant symbionts after a stress, this causes slower coral growth and reduces their recovery. The net effect is that tolerant-dominance seems to hamper coral growth without conferring any survival advantage in response to stress. Indeed, Fig. SM12 suggests that had the corals never switched to tolerant symbionts they would survive just as long.

### 3.3 Thermal scenario 2: Stable future thermal stress intensity

We consider a scenario in which the intensity of thermal stress events is held constant (set to the normalized value of the 2016 event) using two frequencies of stress events (as in the control), that determine whether there is time for symbiont reversal. Regardless of the frequency of stress events, corals survive the duration of 150 simulated years (300 timesteps), with coral cover fully recovering after each heat stress event in both the 3-year and the 5-year scenario (Fig. 5a and 5b respectively). However, when the simulations are extended to longer timescales (e.g., 300 simulated years), high frequency thermal stress events eventually lead to reef collapse (Fig. 5c), while low frequency ensures indefinite survival of the reef (Fig. 5d). Thus, we observe the opposite outcome compared to the increasing intensity scenario where the case of high frequency stress events was associated with longer survival.

**Figure 5:**
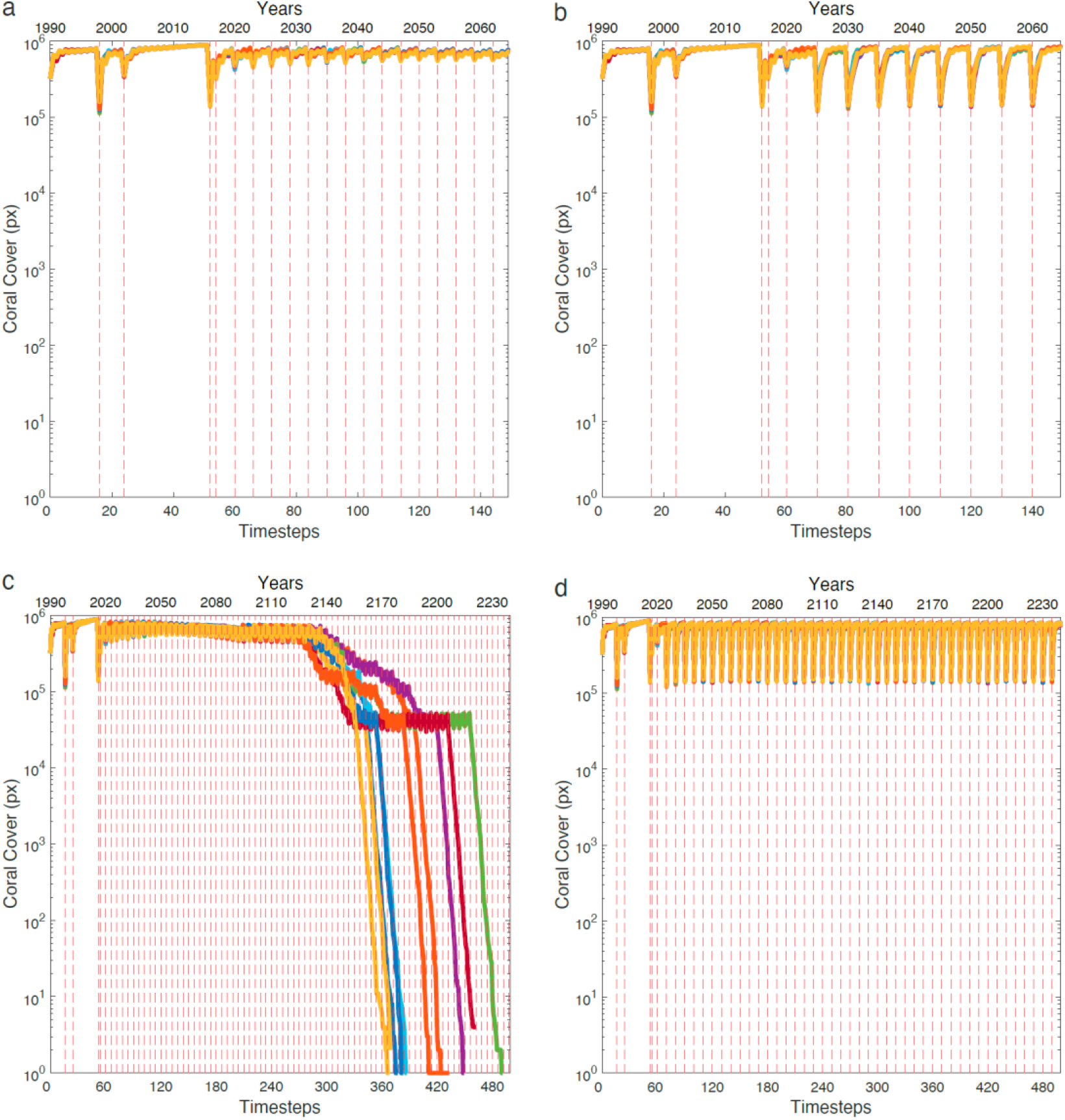
Projected coral cover (as pixels - px) over 75 simulated years (150 timesteps) under thermal stress events of fixed intensity every 3 years (a) or 5 years (b), with deterministic 4-year reversal. In the 3-year cycle (a), thermal stress events occur before thermally-tolerant symbionts can revert to thermally-sensitive symbionts. In the 5-year cycle (b), thermal stress events occur after reversal to sensitive symbionts. Extended simulation to 250 simulated years (500 timesteps) at 3-year (c) and 5-year (d) scenarios are also provided. Ten independent runs are shown for each scenario.

In the stable intensity scenario, low frequency events (5-year cycle) allow indefinite reef survival as, 1) the stress intensity remains within the tolerance limits of sensitive symbionts (see 2016 values in Table SM3), so that even complete sensitive symbiont-dominance at a reef level does not lead to extinction when a stress event occurs. 2) Because there is some time (1 year) between symbiont reversal and the next stress event, corals can take advantage of the faster growth of sensitive symbionts and improve recovery. 3) The probabilistic nature of switching from sensitive to tolerant symbionts ensures that some corals remain thermally sensitive-dominated, which means they can both survive stress events and capitalize on the growth advantage of thermally sensitive symbionts for the entire 5-year cycle. The combination of these effects restores the coral population to pre-disturbance levels after every thermal stress event.

In contrast, when stress events occur at high frequency (3-year cycle), coral cover eventually declines. This occurs because the higher frequency of events causes corals to become dominated by thermally tolerant symbionts. While this offers resistance to thermal stress, it comes at the cost of reduced growth rates in between events. Reef collapse is then driven by the cumulative effect of more frequent disturbances—which reduce the recovery time between stress events and the reduced growth capabilities of thermally tolerant symbiont-dominated corals.

### 3.4 Symbiont populations shape reef structure

The dominance of thermally tolerant symbionts causes corals to grow more slowly during recovery, resulting in eventual reef extinction in the scenario with fixed intensity, high frequency events. To understand how this occurs, we compare the evolution of the reef’s structure by investigating the organization of the reef in corals of different ontogenic stage (and thus size) (Fig. 6). In the scenario of fixed thermal stress intensity, the reduced growth rate associated with full thermally tolerant symbiont dominance leads to structural changes within the reef. Over time, large adult corals dominate the population, while juvenile and pubescent stages decline. Reefs become dominated by adult corals and larval stages that never develop into adults. When the last adult corals die, there is no coral capable of producing larvae and the reef undergoes an abrupt decline. If we explore at the reef structure in the case of fixed intensity low frequency stress events—where corals survive indefinitely— we see a more balanced population structure with full recovery of juveniles and pubescent stages.

**Figure 6:**
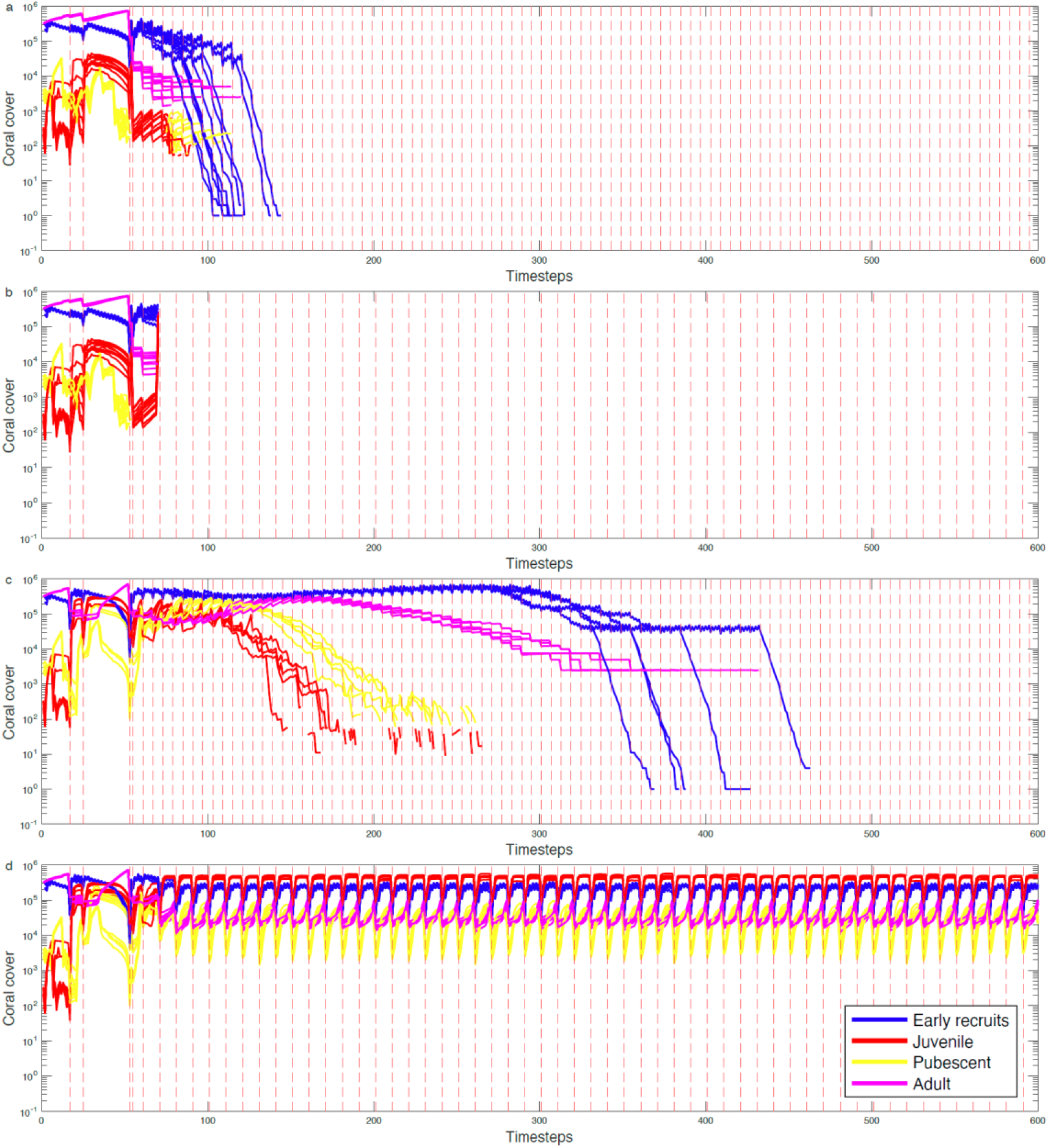
Projected coral cover (as pixels) over time for different life stages under different thermal stress scenarios, with deterministic 4-year reversal. (a) 3-year cycle under increasing thermal stress intensity. (b) 5-year cycle under increasing thermal stress intensity. (c) 3-year cycle under fixed thermal stress intensity. (d) 5-year cycle under fixed thermal stress intensity. Ten independent runs are shown for each scenario.

### 3.5 Interaction between the model and real-world climate projections

Our hypothetical *in silico* simulations show that the characteristics of the stress (intensity and frequency) determine the outcome of stress-induced switching (leading to bleaching) between sensitive and thermotolerant symbionts. We explore the future implication of these hypothetical *in silico* observations using a projection of real-world future thermal stress regime from IPCC SSP-126. The qualitative nature of theses real-world thermal stress events broadly resembles the increasing intensity scenario with a high frequency of events. We observe that coral cover remains reasonably high until the mid-2020s, at this point, all simulations show a consistent decline in coral cover to 2035 which is associated with increasing thermal stress (as DHM) during the summers over that period (Fig. 7). These analyses are consistent with our earlier analyses of the increasing intensity scenario, and provide a demonstration of how our model may interact with real-world data to determine the impact of symbiont switching on coral reef survival trajectories.

**Figure 7:**
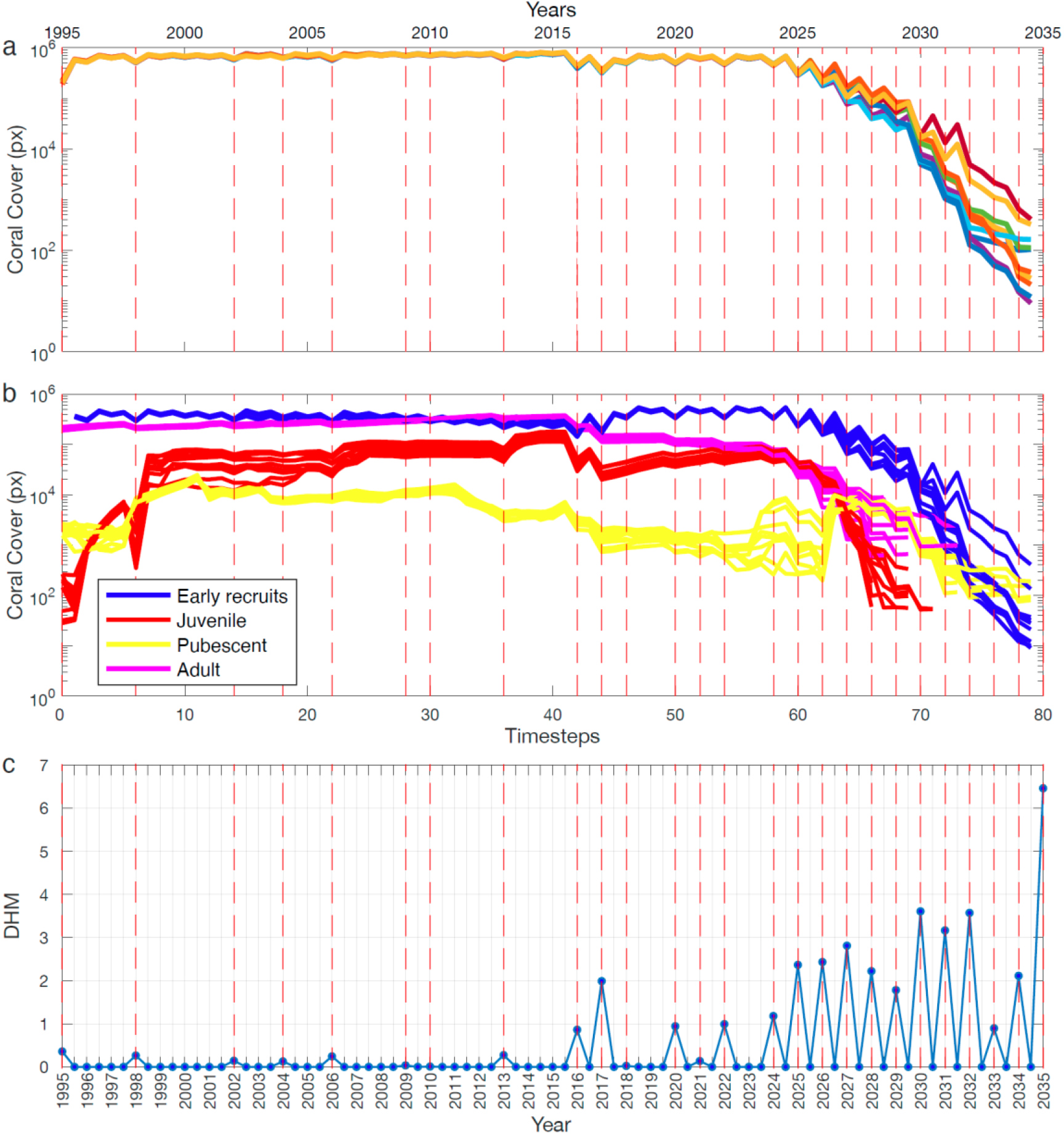
Projected coral cover (as pixels -px) from 1995 to 2035 in our model using real-world historical data (1995–2024) and IPCC SSP-126 projections (2025–2035, ‘low carbon emission’ scenario). Past and future thermal stress is introduced using normalized (to 2016) Degree Heating Months (DHM) as a proxy for coral stress (see SM1.10.4 for full details). a) Total coral cover. b) coral cover separated by life stage. c) Degree heating months. Ten independent runs.

## 4 Discussion

Empirical observations indicate that corals can exhibit increased resilience to heatwaves when these occur in close succession [7], although the mechanisms underlying this pattern remain unclear. Here, we explore those possible mechanisms using forward-pointing models, with both simplified and real-world thermal stress, to investigate the dynamics of coral survival under future warming. We find that the interplay between symbiont switching processes and the timing and intensity of heat stress events determines the long-term survival of coral populations. In particular, we observe that if corals repeatedly switch between thermally-tolerant and thermally-sensitive symbiont populations, this can either ensure long-term survival, acting as an acclimatory process, or reduce their growth to the extent that they are out-competed. Which trajectory occurs depends on the specific temporal and intensity dynamics of thermal stress.

### 4.1 Thermal scenario 1: Increasing future thermal stress intensity

When thermal stress events increase in intensity over time, we find that coral populations ultimately die regardless of whether they harbour thermally sensitive symbionts or ones that endow thermal tolerance. While this initially suggests any future acclimatory advantages corals attain from thermotolerant symbionts may be irrelevant, the amount of time and stress that corals can survive does change depending on the proportion of thermotolerant symbionts they host; i.e., there is scope for a degree of acclimation-driven survival but it may be time limited. For example, we find that corals that experienced higher frequencies of thermal stress survive for longer timescales than those exposed to less frequent thermal stress. This finding does not support the current expectation that higher frequencies of bleaching stress are deleterious but rather supports the acclimatory hypothesis [e.g. 11]. Our model suggests that mechanistically this occurs because when thermal stress is more frequent, it causes surviving corals to continuously harbour a preponderance of thermally tolerant symbionts, allowing them to better survive future thermal stress.

In contrast, when thermal stress events are less frequent, corals switch back to a preponderance of thermally sensitive symbionts [e.g. 16] so their growth is faster [32] but this leaves them susceptible to subsequent thermal stress events [33]. Thus, given that thermotolerant symbionts are generally associated with lower coral growth via reduced calcification (e.g., *Durusdinium* sp [34]) we observe a somewhat counterintuitive outcome in a warming world: despite the sub-optimal growth and reduced competitive advantage, corals survive for longer when continually hosting a preponderance of thermally-tolerant symbionts — a process stimulated by frequent thermal stress events of survivable intensity. While at first, this appears in contrast to some observational studies [e.g. 33] we attribute the difference to the timescales under consideration; experimental approaches generally focus on annual to sub-decadal dynamics and modelling approaches allow us to explore decadal and centennial timescales.

### 4.2 Thermal scenario 2: Stable future thermal stress intensity

Under stable future thermal stress, if stress intensity remains within the tolerance limits of both types of symbionts, coral populations can survive indefinitely by repeatedly switching between symbiont populations. This occurs when stress events repeat at a lower frequency (longer than the symbiont-type reversal time) enabling reversal to a predominance of thermally-sensitive symbionts. The interim period between reversal and the next thermal event allows corals to take advantage of the faster growth of sensitive symbionts to aid reef recovery. Moreover, when thermal stress is stable a small fraction of thermally sensitive symbiont-dominated corals survive heatwaves further promoting reef growth for the entire cycle. However, once bleaching frequencies increase (shorter than the symbiont-type reversal time) thermally tolerant symbionts become exclusively dominant and — even under stable future temperatures — coral cover eventually declines at centennial timescales (ca. 150 years in our simulations, see Fig. 5c). This decline may be due to the reduction of growth [e.g. 35] which causes a disruption of the coral’s turnover cycle and the reef’s age structure (Fig. 6). Both of these disruptions can be generated by the reduced energetic flow observed from thermally tolerant symbionts to their host [2].

Critically, this suggests that even if warming were to be maintained at current temperature levels (sub-lethal for both symbiont types) but that frequencies were to increase, the initial acclimatory-linked survival provided by thermally tolerant symbionts would eventually be overtaken by the negative impacts of a chronically-reduced supply of energy from the symbionts to the coral hosts.

### 4.3 Coral survival under real-world temperature projections

We used IPCC scenarios to provide environmental realism and real-world coral survival trajectories in a future warming world. Even under a low emission scenario, our projections indicate that corals will experience thermal stress above the survival threshold for thermally sensitive symbionts most summers to at least 2035 (the length of our simulations). Although winters do offer important seasonal reprieve from thermal stress, the year-on-year thermal stress during summer eventually results in a notable decline in coral cover in our model, beginning in the mid-2020s, with widespread coral mortality occurring by the mid-2030s. These predictions assume that rising temperatures correspond directly to corals experiencing increased thermal stress. Should corals adapt to temperatures in some way, perhaps via symbiont evolution, then future thermal stress may manifest more like the fixed intensity scenarios we consider. In absence of this adaptation, the projected declines in coral cover trajectories stem from a combination of factors. As thermal stress will have near-annual frequency in the future, with intensity exceeding the tolerance of sensitive symbionts, corals soon become dominated by thermally-tolerant symbionts, hence their growth is slower. As with our simplified stress models, such sustained growth reduction likely contributes to the population’s altered demographic structure such that juveniles and pubescent stages are unable to effectively replace ageing adults (see Fig. 7b). Additionally, thermal stress events remain sufficiently intense to cause elevated mortality even in corals hosting thermally-tolerant symbionts, further weakening the reef over time. By the late 2020s, corals are locked into a cycle of slow growth, reduced recruitment, and ongoing mortalities— culminating in a collapse around the mid-2030s even in the lowest emission IPCC scenarios. Thus, despite any acclimatory benefits offered by hosting thermally-tolerant symbionts, the high frequency of stress events implies that corals cannot sustain high enough growth to ensure future reef survival.

### 4.4 Strategies for future coral survival trajectories

Both our simplified model scenarios and our IPCC-based simulations indicate that under increasing frequency corals tend to host only thermally-tolerant symbionts. Under high thermal intensity, this confers acclimatory advantages at decadal timescales, but these advantages are time-limited and cannot ensure coral survival in the longer term. The individual mechanisms involved in such responses are supported by documented biological processes in real-world reefs. Mortality due to acute heat stress itself is one of the most documented outcomes of recent bleaching events, particularly on the Great Barrier Reef [36, 37], but so is the ability to survive successive, temporally-close, low intensity bleaching events [8, 38]. Separately, naturally slower-growing corals (e.g., massive) are generally outcompeted by faster growing competitors (e.g., branching) as they compete for access to light but ultimately exist on reefs by using niches which are sub-optimal for faster growing corals [39]. Further, temporary slower growth induced by the presence of symbionts endowing thermal tolerance is now recognised as a negative impact of attaining thermal tolerance such that corals migrate back to hosting predominantly symbionts endowing higher growth but lower thermal tolerance [e.g. 18, 40].

However, we now show that all the processes interact to produce different coral survival trajectories driven by the dynamics of their relationships with the frequency and intensity of thermal stress. Critically, contrary to expectations, we show that corals can use two different acclimatory strategies to survive. If thermal stress is low in intensity and frequency then alternating between hosting predominantly thermally-sensitive and thermally-tolerant symbionts causes corals to cycle between high growth and increased thermal resilience in order to remain competitive. If thermal stress is high in intensity and frequency but remains sub-lethal for thermally tolerant symbionts, then continuously maintaining predominantly thermally tolerant symbionts allows corals to acclimate to their warming environment, even while growing more slowly and for a limited time. In both cases, chronic levels of stress (e.g. as generated by low energy flow from the symbionts to the host) may lead to coral reef decline due to changes in coral population structure — an indirect effect of warming oceans.

### 4.5 Future modelling directions

Our model makes advances over previous forward pointing models by also incorporating symbiont reversal in addition to the symbiont switching present previously in Ortiz et al. [21]. This enabled us to explore both the mechanisms by which corals may acclimate to a warming world and the different survival strategies they may use. Our model is simplified by removing some ecological dynamics which allows us to assess generalised future trajectories of coral survival but not finer detail dynamics (e.g. detection of specific years of mass coral death or survival). Such knowledge is beneficial for making informed *a priori* management decisions rather than responding at shorter time scales (e.g., seasonal projections of coral health). We believe this can be achieved by broadening our model to also incorporate empirical physiological data with respect to spatial and species-specific differences in the symbiont communities found within coral hosts. This will produce less parsimonious and more computationally demanding models, but they could still be deployed for specific regions with modest time and financial resources. In addition, we do not consider symbiont evolution in our model, indeed it could be that the symbionts evolve greater thermotolerance, or improved energy transfer to the host leading to different trade-offs for coral growth. As such, the addition of holobiont evolution would allow for not only acclimation responses but also adaptive evolutionary responses, particularly those of the symbiont community, which could feasibly evolve at the centennial timescales we consider here. At the ecosystem level, grazing intensity in our model is fixed which means our models may under-or overestimate macroalgal competition and its negative feedback on coral reproduction when coral cover is low. This does produce agile models deployable in many locations, but for areas with specific or unusual grazing dynamics, we suggest the addition of grazing dynamics and their interplay with the projected warming could help further refine higher spatio-temporal resolution projections of coral health.

## 5 Conclusions

We model coral survival in a future world characterized by stable or warming temperatures where the intensity and frequency of thermal stress events, as well as symbiont switching dynamics, play crucial roles in determining coral survival trajectories. Contrary to expectations we show that corals may use two strategies for future survival, both of which represent coral acclimatory capacity as a means of survival. If thermal stress is low in frequency and intensity, corals alternate between hosting predominantly thermally-sensitive and tolerant symbionts, but alternatively, if thermal stress increases and is frequent, corals maintain predominantly thermally tolerant symbionts. These strategies are not mutually exclusive, and corals may switch between them depending on prevailing environmental conditions. However, using real-world temperature projections, we show that this flexible strategy may still lead to an immediate and sustained fall in coral cover. Critically, by incorporating symbiont thermal sensitivities and reversible switching under different thermal stress frequency and intensity scenarios, our models now provide a realistic yet parsimonious approach for exploring and projecting coral survival trajectories in a changing world.

## 6 Acknowledgments

We thank our IceLab and Departmental colleagues and collaborators for constructive discussions around our research question and manuscript.

