## Supplementary Information for "The dynamics of thermal stress events determine whether thermotolerant endosymbionts improve or hamper survival of coral reefs"

### SM1 Detailed Model Description

Our mechanistic reef model constitutes an independent implementation of a previous model parameterized to represent a Caribbean coral reef ecosystem, originally introduced by Mumby (2006), further refined by Mumby, Hastings, and Edwards (2007), and later expanded to include symbiont switching by Ortiz, Gómez-Cabrera, and Hoegh-Guldberg (2013). While our code was developed independently, most model components follow the mechanistic formulations proposed in these reference works. In the following sections we present our model implementation in parallel with the original formulations, indicating where we have fully adopted the structure and parameters from the reference models, introduced modifications to parameters or dynamics, or implemented entirely new components that were not present in the original frameworks. A concise comparison of key structural and ecological assumptions adopted in different versions of the reef model is reported in Table SM4, located at the end of the Supplementary Material.

#### SM1.1 Macroalgal (MA) growth over corals

##### SM1.1.1 MA overgrowth in Mumby (2006)

In Mumby (2006) macroalgal overgrowth is modeled as the probability that a coral colony is extirpated due to encroachment by mature macroalgae (age > 6 months). At each 6-month timestep, the probability of overgrowth for a coral within a 0.25 $\text{m}^2$ cell is given by:

$$P_{C\to M} = P_{\text{size}} \cdot M_{12} \tag{1}$$

where $P_{C\to M}$ is the probability of extirpation, $M_{12}$ is the proportion of the cell area (2500 $\text{cm}^2$) covered by mature macroalgae, and $P_{\text{size}}$ is a size-dependent vulnerability function:

$$P_{\text{size}} = 0.83 \cdot \exp(-k \cdot x) \tag{2}$$

with $x$ being the coral area (cm$^2$) and $k = 0.0012$, fitted to empirical data from Jamaica. This formulation reflects the higher susceptibility of small corals to overgrowth. For example, a 60 cm$^2$ coral in a cell with 40% macroalgal cover has a 39% chance of extirpation:

$$P_{C\to M} = 0.83 \cdot e^{-0.0012\cdot 60} \cdot 0.4 \approx 0.391$$

These equations are described in detail in the Appendix of Mumby (2006).

##### SM1.1.2 MA overgrowth in Mumby et al. (2007)

In Mumby et al. (2007) direct overgrowth of coral by macroalgae $O_{C\to M}$ is calculated as:

$$O_{C\to M} = M_{4cells} \cdot P_i \cdot 4/7$$

where $M_{4cells}$ is the proportion of macroalgae in the von Neumann (VN) 4-cell neighborhood (which includes the central cell), $P_i$ is the perimeter of the coral, and 4/7 is a scaling factor introduced to take in account experimental data. A complete parametric description, including experimental data, is given in Mumby et al. (2007), SM, Section *Competition between corals and macroalgae 1: effect of macroalgae on corals.*

##### SM1.1.3 MA overgrowth in our model

Macroalgal overgrowth parametrisation differs between Mumby (2006) and Mumby et al. (2007). In our model we parametrise MA overgrowth as in Mumby et al. (2007). Note that MA overgrowth can potentially lead to the extirpation of corals of any size.

#### SM1.2 Coral growth

##### SM1.2.1 Coral growth in Mumby et al. (2007)

In Mumby et al. (2007) "coral size is quantified as the cross-sectional, basal area of a hemispherical colony ($cm^2$). BC (brooding corals) have a lateral extension rate of 0.8 cm yr$^{-1}$ (i.e., 0.4 cm per semester) and SC (spawning corals) grow slightly faster at 0.9 cm yr$^{-1}$ (i.e., 0.45 cm per semester). Calculations are based on median rates for *Porites astreoides*, *P. porites*, *Siderastrea siderea*, *Montastraea annularis*, *Colpophyllia natans* and *Agaricia agaricites*)". Furthermore, direct competition between macroalgae and cover can reduce corals' growth rate:

1. Growth rate of juvenile corals (area $< 60cm^2$) is set to zero if $M_{4cells} > 80\%$, and reduced by 70% (i.e., reduced to 30%) if $60\% < M_{4cells} \leq 80\%$. Parameters based on both Dictyota and Lobophora.

2. Growth rate of pubescent and adult corals (area $\geq 60cm^2$) reduced by 50% if $M_{4cells} > 60\%$

##### SM1.2.2 Coral growth in our model

First of all we calculate a basal lateral extension rate (henceforth refferred to simply as *basal growth rate*) $G_r^b$ for all corals in a given cell of the lattice. This 6-months-basis growth rate depends on the relative quantity between symbionts in each coral: in our paper full SeS occupation endows 0.9 basal growth rate, full ToS occupation endows 0.1 basal growth rate, intermediate occupation gives an interpolated value. Note that these values are not taken from experimental data or literature, but instead used to extremise ToS and SeS features and better discern the differential dynamics of the system. After defining $G_r^b$, we calculate growth reduction due to VN

4-cell neighborhood macroalgae percentage (including the current, central cell), and therefore the effective growth rate $G_r^e$ according to the following scheme:

- if coral is juvenile and $M_{4cells} \geq 80\% \rightarrow G_r^e = G_r^b \cdot 0$ (full inhibition)
- if coral is juvenile and $60\% \leq M_{4cells} < 80\% \rightarrow G_r^e = G_r^b \cdot 0.3$ (partial inhibition)
- if coral is juvenile and $M_{4cells} < 60\% \rightarrow G_r^e = G_r^b$ (no inhibition)
- if coral is pubescent or adult and $M_{4cells} \geq 60\% \rightarrow G_r^e = G_r^b \cdot 0.5$ (partial inhibition)
- if coral is pubescent or adult and $M_{4cells} < 60\% \rightarrow G_r^e = G_r^b$ (no inhibition)

Finally, we calculate growth for each coral in the cell. The growth increment is translated into a required additional area for each coral, and the sum of required growth areas for all colonies in the cell is compared to the available cropped algae (turf) substrate. If the turf is sufficient, all corals grow fully, and turf is proportionally reduced. If turf availability is insufficient, the model prioritizes the growth of the largest coral in the cell. If there is enough turf for the biggest coral to grow fully, then it will do so. If the turf is not enough for the full growth of the biggest coral in cell, then our model allows for two alternative strategies:

- Exclusive use: the largest coral grows only up to the available turf area.
- Competitive exclusion: the largest coral grows fully at the expense of smaller colonies. These may be partially reduced or entirely removed depending on the space needed to grow. If necessary, multiple smaller colonies are removed until the required area is achieved. If no turf is available and multiple corals are present, one smaller coral is removed to allow full or partial growth of the largest. In the paper's results we always follow the competitive exclusion strategy.

If only one coral is present and no turf remains, growth does not occur as a single coral cannot exceed the cell's boundaries.

#### SM1.3 Vegetative macroalgal growth

In Mumby (2006) and Mumby et al. (2007) macroalgae can arise either from aging of cropped algae that are not grazed (see Section SM1.5), or from vegetative growth over cropped algae. From Mumby et al. (2007), SM: "The probability that macroalgae will encroach onto the algal turf within a cell, $P_{CA \rightarrow MA}$, is given by:

$$P_{CA \rightarrow MA} = M_{4cells}$$

where $M_{4cells}$ is the percent cover of macroalgae within the VN 4-cell neighborhood. This is a key method of algal expansion and represents the opportunistic overgrowth of coral that was extirpated by disturbance." Moreover, due to competition between corals and MA, $P_{CA \rightarrow MA}$ is reduced by 25% when at least 50% of the local VN neighborhood includes coral, so that:

$$P_{CA \rightarrow MA} = 0.75 \times M_{4cells} \text{ if } C \geq 0.5$$
$$P_{CA \rightarrow MA} = M_{4cells} \text{ if } C < 0.5$$

In our paper we implement vegetative MA growth exactly as in Mumby et al. (2007). For references to literature see both Mumby (2006)(Appendix) and Mumby et al. (2007)(SM, sections *'Macroalgal growth over dead coral (cropped algae)'* and *'Competition between corals and macroalgae 2: effect of corals on macroalgae'*).

#### SM1.4 Grazing

##### SM1.4.1 Grazing in Mumby (2006)

Grazing in the model is driven by herbivorous parrotfishes (scarids) and, in some scenarios, by the sea urchin *Diadema antillarum*. Grazers remove algae from the reef, maintaining substrate in a cropped algal state and preventing macroalgal overgrowth. Grazing occurs every six months, with cells selected in random order. Mainly only parrotfish grazing is considered, as it is assumed in Mumby (2006) that sea urchin is depleted in the Caribbean. It is considered in the model that parrotfish do not increase total grazing in response of increased food (i.e., algal) availability. Therefore grazing levels are stable and independent of coral cover levels. Although this is not the case for urchin grazing (Mumby (2006), [Levitan, D. R. 1988b. Density-dependent size regulation and negative growth in the sea urchin Diadema antillarum Philippi. Oecologia 76:627–629.]), this is also fixed in the model when introduced.

Parrotfish grazing is spatially constrained: each timestep allows a fixed proportion of the reef to be grazed, based on the assumed biomass of the fish community. Three scenarios are defined: low (10%), medium (20%), and high (30%) grazing, representing increasing levels of fish biomass and reef protection. Once the grazing threshold is reached, no additional algae are removed during that timestep. Grazers do not discriminate between cropped algae and macroalgae, and coral recruits are not grazed. Cells are grazed in a random order and all cropped algae and macroalgae are converted to (or remain in) the initial cropped algal state until the spatial constraint is reached (e.g., up to 30% of the total reef area is successfully grazed every 6 months). Increase or decrease of parrotfish population due to higher or lower algal biomass availability is not taken in account.

Urchin grazing is modeled independently, assuming *Diadema* can graze up to 40% of the reef surface when present, based on historical removal rates. Parrotfish and urchins graze in overlapping spatial distributions, meaning that the same CA or MA patch can be grazed independently by both grazers and that the same patch adds up independently to the maximum grazing percentage per timestep for each grazer type.

##### SM1.4.2 Grazing in Mumby et al. (2007)

In Mumby et al. (2007) the main mechanism is mostly the same as in Mumby et al. (2007), but the percentages change slightly. An unfished community of parrotfishes can graze a maximum of 40% of the seabed per 6 month time interval. Urchin, when present (in the majority of simulations they are considered depleted) can graze up to 53% of the seabed. When both grazers are present, there is an overlap between areas grazed by parrotfish and urchin, to that the total grazed area every 6 months is less than the sum of the two maximum grazable areas. Finally, in Mumby et al. (2007) parrotfish can predate a maximum 15% of recruits for each 6 month iteration, resulting in dead coral (i.e., cropped algae, as there is no distinction between the two in this model). Parrotfish recruits predation is confined to small corals of area $\leq 5cm^2$.

##### SM1.4.3 Grazing in our model

Grazing in our model has the same general characteristics as Mumby et al. (2007), with the difference that only one generic grazer is considered, with a grazing capability of 80% of the seabed per 6 month time interval. With respect to recruit grazing our generic grazer behaves substantially as parrotfish and can predate a maximum 15% of recruits ($\leq 5cm^2$) for each 6 month iteration, resulting in dead coral/cropped algae.

#### SM1.5 Aging of macroalgae and cropped algae

##### SM1.5.1 Algal aging in our model

Algae are modeled as two functional groups: cropped algae (CA) and macroalgae (MA), each tracked by age class within 0.25 $m^2$ cells. Macroalgae become cropped algae when grazed. Cropped algae have 5% probability of becoming macroalgae in the same semester of grazing, and 100% probability after one semester without grazing. Therefore, compared to (Mumby, 2006) and (Mumby et al., 2007), the onset of macroalgae occurs more rapidly. This is compensated by the higher grazing levels.

#### SM1.6 Larval recruitment

##### SM1.6.1 Larval Recruitment in Mumby (2006)

The specific details on how coral recruitment is modeled in Mumby (2006) are unclear. In the main paper it is declared that two different reproductive modes are implemented, where brooders (e.g., *Porites astreoides*) produce locally retained larvae, while spawners (e.g., *Siderastrea siderea*) release gametes that disperse regionally. However, when dispersal among connected reefs is described in the rest of the paper (4 reefs are connected with unidirectional currents transporting larvae from one reef to the next), only retention and dispersal coefficients are indicated, but it is not clear if dispersed larvae belong only to spawners and retained ones only to brooders. Recruitment is implemented every six months, with coral settlers assigned to cropped algae substrata (i.e., turf). Corals recruit into the model at a density of 2 individuals every $0.25m^2$ (or one cell) of cropped algae. This implies a probability of successful recruitment of $P_r = 2$ recruits/$2500cm^2$ of CA $= 0.0008$ recruits per $cm^2$ of CA (1 pixel = 1 $cm^2$). Fertility is size-dependent, with pubescent corals ($60 - 250cm^2$) exhibiting 25% fecundity compared to adult ($> 250cm^2$) corals. It is also indicated that larval production is modeled on Porites astreoides, which has approximately two eggs per gonad, six gonads per polyp, and 18 polyps per $cm^2$ of coral (Szmant, 1986). It is however unclear whether these biological values are used in the model, and if so, how. Finally, a set of equations is given to indicate the probability of successful recruitment $P_r$ depending on whether a stock-recruitment relationship (recruitment is proportional to fertile corals' area) or full open recruitment (recruitment at max values) is implemented:

$$P_r = 1 \quad \text{if} \quad C \geq 30\% \quad (T_1 \approx 2.9 \times 10^9)$$
$$P_r = 0 \quad \text{if} \quad C \leq 3\% \; (T_1 \approx 2.9 \times 10^7)$$
$$0 < P_r < 1 \quad \text{if} \quad 3\% \leq C \leq 30\%$$

where $T_1$ represents total larval input for various stock sizes. It is however unclear how the $T_1$ values are calculated and how probabilities of recruitment are defined. Speculating, if one were to calculate larval production in a $50 \times 50$ reef composed of cells of $2500cm^2$ each, using the biological values indicated above, one would get $\approx 4 \times 10^8$ larvae for 30% coral cover, and $\approx 4 \times 10^7$ larvae for 3% coral cover.

##### SM1.6.2 Larval Recruitment in Mumby 2007

In Mumby et al. (2007)(SM) it is indicated that coral recruits are able to recruit on cropped algae, with a recruitment rate of 2 recruits (0.2 recruits) per $0.25m^2$ of cropped algae per time interval per brooders (per spawners). These numbers were adjusted for rugosity (∼2) and the cover of cropped algae at Glovers Reef (Mumby, 1999). In the main paper it is indicated that there is no stock-recruitment relationship and that larvae recruit at maximum levels (up to 4 per $0.25m^2$, probably rugosity is take in account here) irrespective of stock size. In the Supplementary Material it is stated that corals are subjected to size-dependent fecundity, nevertheless it is also stated that reproduction of corals within the reef is excluded, and that a constant rate of coral recruitment from outside reef (i.e. no stock-recruitment dynamics) is assumed.

##### SM1.6.3 Larval recruitment in our model

Coral larvae are produced by fertile colonies in adult stage every 12 months (i.e. every two timesteps). Larval output is proportional to colony area and modulated by macroalgal cover in the surrounding $M_{5cell}$ neighborhood: colonies exposed to high macroalgal abundance experience reduced fecundity (25% reduction if $M_{5cell} > 0.5$). Total larval production is defined by the effective fecundity of each coral multiplied by its area. In our model, fecundity = 10, and each larva has a size of $1cm^2$ (i.e., it occupies 1 pixel in the lattice). This means that each coral in the reef can produce up to 10 times more larvae than its area (if no MA fecundity reduction is considered). Therefore a limited number of corals is already able to saturate the recruiting need of the reef, and when many corals are present more larvae are produced than the ones that can be accomodated by the reef. Larval production is different from recruiting: recruiting is defined as the successful attempt by larvae of colonising turf. In our model, after applying a fixed pre-recruitment mortality (10%), surviving larvae are randomly assigned to available turf cells, without imposing a strict limit on the maximum number of corals per cell, meaning that virtually an entire cell can be colonised by larvae if enough larvae and turf are available. Each larva inherits key properties from its parent colony, including current symbiont community composition. Recruitment continues iteratively until either all larvae are placed or available turf is exhausted.

#### SM1.7 Corals mortality except thermal stress events

In our model corals can die due to macroalgal overgrowth (any size, see *Macroalgal overgrowth* subsection), thermal stress (any size, see *Thermal stress events* subsection), grazing (only early recruits, $\leq 5cm^2$), random whole colony mortality (only pubescent corals, $60 - 250cm^2$). Note that juvenile ($5-60cm^2$) and adult corals ($\geq 250cm^2$) can only die due to macroalgal overgrowth or thermal stress events. Early recruits mortality due to grazing is set at 15% every 6 months. Random pubescent whole corals mortality is set at 2% every 6 months. Partial mortality and mortality due to hurricane events are not implemented in our model.

#### SM1.8 Corals mortality due to thermal stress events

Unlike the original model in Mumby (2006) and Mumby et al. (2007), which only included coral mortality due to hurricanes, our framework explicitly incorporates thermal stress–induced bleaching mortality, as proposed in Ortiz et al. (2013).

Bleaching-induced coral mortality is activated during periods of thermal stress. At each stress event, the mortality risk of individual corals is calculated as a function of their symbiont composition and the intensity of the heatwave.

For each coral, the coral's sensitivity to thermal stress is calculated as a linear interpolation of the thermal sensitivity associated with each symbiont type, weighted by their relative abundance within the coral. This allows intermediate symbiont compositions to modulate coral sensitivity to stress. Specifically, in our model, thermally sensitive symbionts confer a mortality probability of 0.9, while thermally tolerant symbionts confer a mortality of 0.1 in a fixed intensity scenario (i.e., when thermal stress intensity is equal to 1). A heatwave intensity of 1 therefore corresponds to the reference mortality endowed by the symbionts; values below or above 1 proportionally decrease or increase the mortality risk. If the final computed probability exceeds 1 due to extreme stress or symbiont weighting, it is capped at 1.

For comparison, in Ortiz et al. (2013) the reduction in bleaching mortality given by the thermally tolerant symbiont compared to the thermally sensitive one is set at 30%, meaning that if bleaching mortality is 0.9 for the sensitive symbiont, it will be 0.63 (i.e., a 30% reduction) for the thermally tolerant one.

#### SM1.9 Symbiont switching and reversal

In our model, we simulate thermal stress–induced symbiont switching from a thermally sensitive to a thermally tolerant symbiont type, following a mechanism similar to that described by Ortiz et al. (2013). In addition, we implement symbiont reversal—i.e., the return from a thermally tolerant to a thermally sensitive community—if no further thermal stress events occur over a predefined period.

Symbiont switching allows coral colonies to replace their dominant symbiont community with an alternative community better suited to elevated temperatures when thermal stress occurs. Switching is probabilistic and modulated by thermal stress intensity. Corals can only switch if they are in a basal symbiont configuration—i.e., if they are dominated by thermally sensitive symbionts and have not already switched in response to a prior heatwave.

The switching probability is set to 0.8 for heatwaves of normalised intensity 1 and scales linearly with heatwave intensity. Thus, more intense heatwaves lead to higher switching probabilities, eventually reaching saturation ($P_{\text{switching}} = 1$). In the same way, less intense heatwaves lead to lower switching probabilities, eventually reaching $P_{\text{switching}} = 0$.

After switching, corals remain in a stressed, thermally tolerant–dominated state for a fixed period (8 semesters in the main paper). Once this reversal period has elapsed—and provided no further heatwaves have occurred—the coral reverts to its basal (i.e., thermally sensitive–dominated) symbiont composition. If an additional heatwave occurs while the coral is still in its stressed state, the reversal is postponed by another 8 semesters.

The reversal process used in the main paper is *deterministic* and *abrupt*, meaning that corals are stably dominated by thermally tolerant symbionts with a fixed composition (90% ToS, 10% SeS) throughout the stressed period (i.e., for 8 semesters after the heatwave), and then revert abruptly to a thermally sensitive–dominated composition (10% ToS, 90% SeS) after exactly 8 semesters if no additional heat stress occurs.

All results shown in the main paper refer to this abrupt deterministic reversal. In the Supplementary Material, we explore two alternative reversal dynamics: abrupt but *probabilistic* reversal, and deterministic but *gradual* reversal (i.e., symbiont composition gradually reverts in a linear fashion).

This symbiont switching and reversal implementation supports flexible and ecologically realistic dynamics of symbiont adaptation, combining stochasticity, thermal dependence, and memory-like reversal delays. Such dynamics may offer a mechanistic explanation for empirical observations of increased resilience of corals to repeated, closely spaced heatwaves—such as those reported by Hughes, Kerry, Connolly, Álvarez Romero, Eakin, Heron, Gonzalez, and Moneghetti (2021)—via transient and reversible shifts in symbiont composition.

#### SM1.10 Construction of the Thermal Stress Sequence

To simulate historically realistic thermal stress scenarios, we constructed a synthetic sequence of heat stress events inspired by empirical observations. Our approach builds upon three steps:

(i) extraction and normalization of historical heatwave data from five major bleaching events on Low Isles (GBR) as reconstructed by Kamenos and Hennige (2024), (ii) artificial extension of the historical sequence simulating either a fixed or a constantly increasing thermal stress intensity at a constant pace (3 or 5-year period) for future years, and (iii) merging of historical data (OISST dataset Huang, Liu, Banzon, Freeman, Graham, Hankins, Smith, and Zhang (2020)) and future projections (IPCC SSP-126). For the simulations in the main paper we used a combination of (i) and (ii), except for the results of Section 3.5 (*Interaction between the model and real-world climate projections*) and Fig. 8, where we used (iii). In Section SM2 we also explore results obtained using a modified version of (ii) only, i.e., where we start from null intensity and increase thermal stress in a linear fashion at a constant pace (3 or 5-year period), with no initial historical sequence.

##### SM1.10.1 Extraction and normalization of historical heatwave data

For the historical sequence, we extracted observed historical heat stress values (Degree Heating Weeks, DHW) from (Kamenos and Hennige, 2024) for five representative heatwave events (1998, 2002, 2016, 2017, 2020) in Low Isles location on the Great Barrier Reef (16.375 oS 145.575 oE), as summarized in Table SM1. The median value (2016, 2.62 maximum daily DHW) was then used as a normalization reference, i.e., it was normalized to 1, and the maximum daily DHW values for the other years were rescaled accordingly. The normalized value of 1 corresponds to a thermal stress intensity where the corals' response is nominal, i.e., where symbiont switching probability is 80%, and where mortality probability is 90% when harboring thermally sensitive and 10% when harboring thermally tolerant symbionts. Since these values depend on thermal stress intensity, coral response is rescaled accordingly when thermal stress intensity changes.

| Year | Max daily DHW (Low Isles) | Normalized (Median-based) stress intensity |
|---|---|---|
| 1998 | 0.31 | 0.12 |
| 2002 | 1.16 | 0.44 |
| 2016 | 2.62 | 1.00 |
| 2017 | 11.50 | 4.39 |
| 2020 | 5.54 | 2.11 |

Table SM1: Historical maximum daily Degree Heating Week values extracted from observational records. The first column contains the year of five major bleaching events as observed in Low Isles. The second column contains the maximum daily DHW values for Low Isles as calculated by (Kamenos and Hennige, 2024). The third column contains stress intensity values normalized using the median event value (2016, 2.62 DHW) as reference value.

##### SM1.10.2 Construction of post-2020 thermal stress sequence

After priming our simulations with historical data, we extend thermal stress sequence in a more controlled and predictable manner, to allow an easier analysis of our system's dynamic behaviour. We can extend the sequence either with fixed intensity heatwaves, or with increasing intensity heatwaves. In both cases the spacing between events is kept at a constant pace (i.e. every 3 or 5 years). In the fixed intensity scenarios, all thermal stress events after 2020 occur with the same intensity of the 2016 event (2.62 DHW), normalized to 1. For the increasing intensity scenarios, normalized thermal stress intensities were extended incrementally. Starting from the 2020 value of 2.11 normalized units ($n.u.$), each subsequent event was increased by $+0.1$ $n.u.$ per event (except a +0.09 $n.u.$ increase for the 2021 event so that its value is 2.2 $n.u.$), simulating a gradual temperature increase (see Table SM2).

Table SM2: Example of projected future heat stress increments.

| Synthetic Year | Normalized Stress Intensity |
|---|---|
| 1998 | 0.12 |
| 2002 | 0.44 |
| 2016 | 1.00 |
| 2017 | 4.39 |
| 2020 | 2.11 |
| 2020 + 1 stress event | 2.20 |
| 2020 + 2 stress events | 2.30 |
| 2020 + 3 stress events | 2.40 |
| ⋮ | ⋮ |
| 2020 + $n$ stress events | $2.10 + 0.1 \times n$ |

Two frequency scenarios were implemented with heatwaves recurring every 3 or 5 years. The incremental increases in intensity were kept identical across both scenarios to ensure comparable long-term cumulative thermal exposure after an equal number of stress events. This means however that year-wise (i.e., after a given amount of time from the beginning of the simulations) corals in the 3-year cycle scenario will have experienced not only more thermal stress events but also higher thermal stress intensity compared to the 5-year cycle scenario (see Fig. SM1a).

###### SM1.10.3 Effects of varying heatwave intensity on symbiont-dependent dynamics

As outlined in Section SM1.9, symbiont switching probability increases linearly with thermal stress intensity, with a baseline probability of 0.8 when thermal stress intensity equals 1 normalized unit ($n.u.$). For intensities exceeding 1.25 $n.u.$, switching probability saturates at 1. The switching dynamics are summarized in Table SM3, column 4, and in Fig. SM1c.

Coral mortality probability after a thermal stress event depends on: (i) symbiont composition (sensitive or tolerant), (ii) the symbiont-specific nominal value (i.e., at intensity 1) for $P_{\text{mortality}}$, and (iii) the thermal stress intensity. In all our simulations, thermally sensitive symbionts have a nominal mortality probability of $P_{\text{mortality}}(\text{SeS}) = 90\%$, while thermally tolerant symbionts have $P_{\text{mortality}}(\text{ToS}) = 10\%$. Columns 5 and 6 in Table SM3 summarize $P_{\text{mortality}}$ values at different intensities for full symbiont dominance. Note that these values assume that corals are fully dominated by either sensitive or tolerant symbionts, while in our simulations the dominant symbiont never exceeds 90% of a coral's internal composition.

Since coral mortality probability increases or decreases proportionally with thermal stress intensity, these values can reach saturation (i.e., full mortality) at a given thermal stress threshold. For fully SeS-dominated corals, the thermal tolerance threshold is set at 1.1 $n.u.$, while for fully ToS-dominated corals it is set at 10 $n.u.$, which is never attained in our simulations. This means that ToS-dominated corals (even at 90% composition) never reach their thermal tolerance threshold in the results presented in our main paper, implying that ToS-dominated corals always retain a non-zero probability of survival after thermal stress events in our simulations.

This feature of our model is highly relevant for the reef dynamics explored, as ToS-dominance consistently represents a viable strategy for maintaining at least partial reef survival under thermal stress events. Nevertheless, this does not guarantee that a ToS-dominated reef can survive

| Year | Value (DHW) | Normalized value | $P_{SeS \to ToS}$ | $P_{\text{mortality}}$ (100% SeS) | $P_{\text{mortality}}$ (100% ToS) |
|---|---|---|---|---|---|
| 1998 | 0.31 | 0.12 | 0.096 | 0.108 | 0.012 |
| 2002 | 1.16 | 0.44 | 0.35 | 0.396 | 0.044 |
| 2016 | 2.62 | 1 | 0.8 | 0.9 | 0.1 |
| 2017 | 11.5 | 4.39 | $> 1$ (Saturated) | $> 1$ (Saturated) | 0.439 |
| 2020 | 5.54 | 2.11 | $> 1$ | $> 1$ | 0.211 |
| +1 | – | 2.2 | $> 1$ | $> 1$ | 0.22 |
| +2 | – | 2.3 | $> 1$ | $> 1$ | 0.23 |
| +3 | – | 2.4 | $> 1$ | $> 1$ | 0.24 |
| +4 | – | 2.5 | $> 1$ | $> 1$ | 0.25 |
| +5 | – | 2.6 | $> 1$ | $> 1$ | 0.26 |
| +6 | – | 2.7 | $> 1$ | $> 1$ | 0.27 |
| ⋮ | ⋮ | ⋮ | ⋮ | ⋮ | ⋮ |
| END3Y | – | 6.0 | $> 1$ | $> 1$ | 0.6 |
| END5Y | – | 4.4 | $> 1$ | $> 1$ | 0.44 |

Table SM3: Year-by-year values of thermal stress events in the increasing intensity scenario used in the main paper and corresponding coral response parameters. Values from 1998 to 2020 are taken from Kamenos and Hennige (2024) and are normalized to the 2016 median event. The last two rows of the table indicate the final normalized values for the 3- and 5-year scenarios, respectively, with the corresponding response parameters. The column $P_{SeS \to ToS}$ indicates the probability of symbiont switching from thermally sensitive to thermally tolerant symbionts. Columns 5 and 6 report the $P_{\text{mortality}}$ for corals fully dominated by thermally sensitive (SeS) or thermally tolerant (ToS) symbionts. At a normalized intensity of 1 (i.e., the 2016 event, black dashed vertical line), $P_{\text{mortality}}(\text{SeS}) = 0.9$ and $P_{\text{mortality}}(\text{ToS}) = 0.1$. If thermal intensity increases or decreases, these values scale accordingly in a linear fashion. Values in red indicate saturation, i.e., probabilities exceeding thresholds, leading to certain switching or mortality. Saturation never occurs for $P_{\text{mortality}}(\text{ToS})$. Note that all switching and mortality probabilities in this table are calculated for hypothetical complete symbiont dominance, while our simulations constrain maximum symbiont dominance to 90%. Additionally (in gradual reversal dynamics only), symbiont compositions can assume intermediate levels. Therefore, the mortality probabilities shown for fully SeS- or ToS-dominated corals should serve only as reference points to understand general coral behavior in our simulations.

indefinitely in our simulations (and indeed, in our results, it does not), as other dynamics—such as cumulative mortality and reduced growth—also play critical roles.

The dynamics outlined in this section are summarized in Fig. SM1, which shows the timing of thermal stress events, the scaling of coral mortality probabilities, and the increase and saturation of symbiont switching probabilities with rising thermal stress intensity. The figure complements TableSM3 and provides an intuitive overview of how thermal stress shapes symbiont-dependent coral survival in our model.

###### SM1.10.4 Construction of a real-world thermal stress sequence: merging of historical data and future projections

In SM1.10.1 and SM1.10.2 we explained how we built the thermal stress sequence for all the results in our main paper except Section 3.5 and Fig. 8, using a combination of a historical thermal stress sequence to prime our simulations, followed by a more controlled sequence of thermal stress events at fixed or increasing intensity with a constant pace of 3 or 5 years. For Section 3.5 and Fig. 8 we used instead a thermal stress sequence which results from the combination of historical OISST data and future IPCC projections. This involved i) retrieving the necessary dataset from online databases, ii) defining a useful proxy for thermal stress which could be used for both past data and future projections, and finally iii) defining a connection point between past data and future projections so that the two sets could be merged in a meaningful way. For past to present data we used OISST dataset (Huang et al., 2020). This dataset is developed using an optimum interpolation (OI) technique, has a spatial grid resolution of 0.25 degree and offers daily sea surface temperature values from September 1981 to present day. For future data we used IPCC SSP-126 projections.

*Use of Degree Heating Months (DHM) as a proxy for Thermal Stress*

For this particular thermal stress sequence, we quantify the intensity of heat stress events using Degree Heating Months (DHM), a cumulative thermal stress metric inspired by the more commonly used Degree Heating Weeks (DHW) and originally proposed in (Donner, Skirving, Little, Oppenheimer, and Hoegh-Guldberg, 2005; Donner, Knutson, and Oppenheimer, 2007). Both metrics track the accumulation of positive sea surface temperature (SST) anomalies relative to a climatological reference threshold, usually defined as the Maximum Monthly Mean (MMM) plus an additional temperature (e.g., a $28°C$ MMM plus $1°C = 29°C$). However, whereas DHW integrates daily anomalies over a 12-week window, DHM accumulates monthly anomalies over a cumulative stress window of defined duration (e.g., 4-months window). This allows to perform a measure of cumulative stress also for future projections, where temperature data is only available on a monthly basis instead of a daily one. As for DHW, DHMs are calculated accumulating over a given period of time the sum of temperatures only when they surpass a threshold given by the climatology (i.e., the maximum monthly mean or MMM) plus an additional temperature threshold. To give an example, if the MMM for a given month (e.g., May) in a given location on the map is $28°C$, and if the threshold to DHM contribution is $1°C$, then this means that that month will contribute to DHM calculation only if $T > 29°C$. If, for example, MMM temperature in May is $T > 30.5°C$, this means that the contribution of May to total DHM calculation is $T = 2.5°C$. The total stress accumulation is then calculated by summing DHM contribution over a window of several months (e.g. 4 months), so that the DHM at time $t$ is given by:

$$\mathrm{DHM}_t = \sum_{i=t-N+1}^{t} \max(0, \mathrm{SST}_i - \mathrm{MMM} + \mathrm{threshold})$$

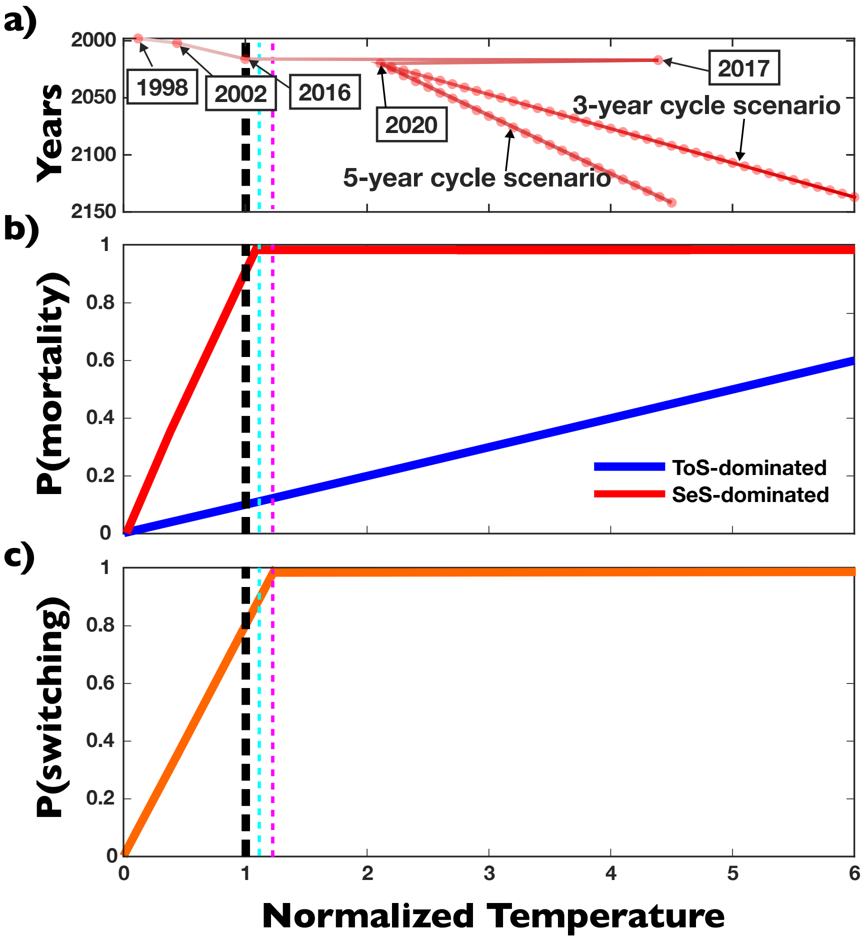


Figure SM1: Relationship between thermal stress intensity, event timing, symbiont switching probability, and coral mortality in the increasing intensity scenario used in the main paper. (a) Schematic of thermal stress intensity variation for the historical sequence (1998, 2002, 2016, 2017, 2020) and for the subsequent projected 3-year and 5-year cycle scenarios. (b) Coral mortality probability ($P_{\text{mortality}}$) as a function of normalized thermal stress intensity for corals fully dominated by thermally sensitive symbionts (SeS, continuous red line) or thermally tolerant symbionts (ToS, continuous blue line). At a normalized intensity of 1, $P_{\text{mortality}}(\text{SeS}) = 0.9$ and $P_{\text{mortality}}(\text{ToS}) = 0.1$. Note that $P_{\text{mortality}}$ scales linearly with thermal intensity until it saturates ($P_{\text{mortality}} = 1$). Saturation occurs at an intensity of approximately 1.1 for SeS-dominated corals, while ToS-dominated corals do not reach saturation for the temperatures attained in our simulations. (c) Symbiont switching probability ($P_{\text{switching}}$) from thermally sensitive to thermally tolerant symbionts as a function of normalized thermal stress intensity. Switching probability increases linearly with thermal stress intensity up to 0.8 at an intensity of 1, saturating ($P_{\text{switching}} = 1$) for intensities exceeding 1.25. The black dashed vertical line indicates the values for normalized temperature, $P_{\text{mortality}}(\text{SeS})$, $P_{\text{mortality}}(\text{ToS})$, and $P_{\text{switching}}$ for the fixed intensity scenario. The cyan and magenta dashed vertical lines indicate the saturation thresholds for $P_{\text{mortality}}(\text{SeS})$ and $P_{\text{switching}}$, respectively. Note that in the fixed intensity scenario, none of the probabilities is saturated, while in the increasing intensity scenario, all probabilities except $P_{\text{mortality}}(\text{ToS})$ (which never saturates in our simulations) are already saturated starting in 2017.

where $N$ is the window size (4 months in our example), and $\text{SST}_i$ is the mean sea surface temperature of month $i$. For our example, with a 4 months window, a $1°C$ threshold, we then have:

$$\text{DHM}_{(\text{May})} = \sum_{i=Feb}^{May} \max(0, \text{SST}_i - \text{MMM}_i + 1°C)$$

This implementation mimics the conceptual structure of DHW as described in NOAA Coral Reef Watch protocols (Liu, Strong, Skirving, and Arzayus, 2006), but shifts the temporal scale from weeks to months as proposed in Donner et al. (2005). Importantly, as per DHW, the DHM value at each timestep serves as a proxy for coral exposure to prolonged heat stress and can trigger bleaching-related dynamics such as mortality or symbiont switching.

The main difference in our implementation compared to (Donner et al., 2005) is the different climatology threshold ($1°C$ in Donner et. al, $.25°C$ in our model) and a slightly different cumulative stress window (4 months in Donner et. al, 3 months in our model).

While DHW remains the operational standard for real-time monitoring, our adoption of DHM allows integrate past sea surface temperature data with future projections. In fact, while the OISST dataset of past sea surface temperatures has a daily resolution, which allows to calculate DHW metrics, the IPCC SST projections have a monthly resolution, which does not allow to calculate DHW. On other other, DHM can be calculated from both datasets and allows the merging of the two in a single thermal stress sequence containing both past and future events

*Merging of historical data and future projections*

To merge historical data and future projections, we selected past observational data ranging from January 1994 to December 2024, and future projections ranging from January 2025 to December 2100. We then converted all data to monthly resolution. This involved converting past data from daily to monthly resolution (by calculating the monthly average temperature), while the future data already had monthly resolution.

We also had to adjust the spatial resolution, as past and future datasets differ in this regard. The coordinates of the geographical area under consideration (Low Isles, Queensland, GBR) are:

- -16.1750 (Northern latitude)
- -16.5750 (Southern latitude)
- 145.3750 (Western longitude)
- 145.7750 (Eastern longitude)

For past data, this subset corresponds to a spatial resolution of $9 \times 9$ pixels, while future projections have a resolution of $2 \times 2$ pixels over the same area. To enable comparison, we upscaled the future dataset to $9 \times 9$ pixels using the *nearest neighbor* interpolation method, assigning to each new pixel the value of the nearest original $2 \times 2$ cell.

We then computed DHM values from 1994 to 2100 for each pixel in our spatial dataset, using the method described earlier in this section and adopting as the baseline climatology the one calculated for the NOAA Coral Reef Watch suite, as described in Heron, Liu, Eakin, Skirving, Muller-Karger, Vega-Rodriguez, De La Cour, Burgess, Strong, Geiger, Guild, and Lynds (2014). Once DHMs were calculated for each pixel and timestep, we selected the maximum DHM value across the spatial subset for each month—yielding a single representative (maximum) DHM value for the entire geographical region from January 1994 to December 2100.

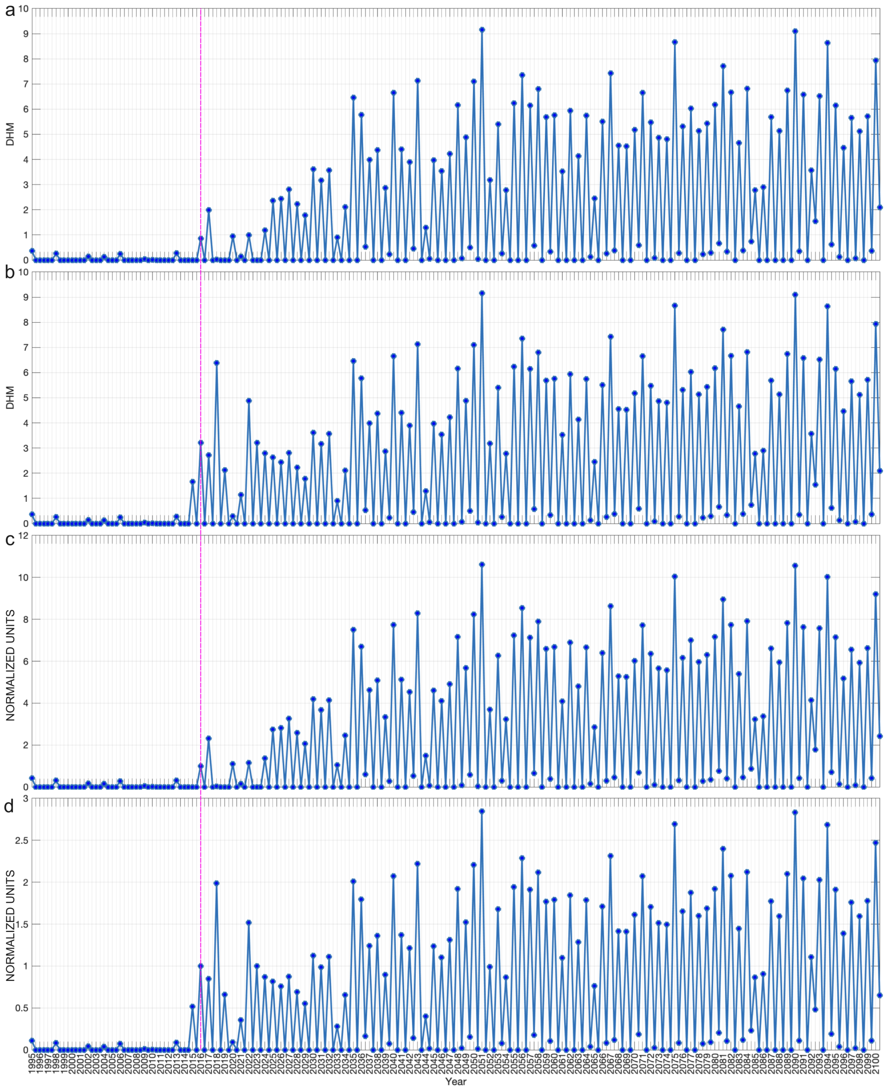


Figure SM2: a) Maximum DHM value for each semester, merging cut-off January 2025; b) Maximum DHM value for each semester, merging cut-off January 2015; c) Maximum DHM value for each semester (normalised to 1st semester 2016), merging cut-off January 2025; d) Maximum DHM value for each semester (normalised to 1st semester 2016), merging cut-off January 2015. For all images we use a climatology threshold of $0.50°C$ and a cumulative stress window of 3 months. The dashed vertical line in magenta indicates the 2016 1st semester peak, which is used for normalization. In a) the 2015-2025 range is drawn from historical series, while in b) it is drawn from the future projections dataset. Similarly, in c) the entire thermal stress sequence is normalised to the 2016 (1st semester) historical value, while in d) it is normalised to the 2016 (1st semester) value drawn from the future projections dataset.

Next, we merged the two datasets. Since the historical dataset ends in December 2024 and the future projections start in January 2015, we had to determine the appropriate cut-off point. We noted that in the overlapping 2015–2024 period, the datasets show a high level of discrepancy—at least in the geographical region considered—with historical data displaying considerably lower thermal intensity peaks compared to future projections (cf. Figs. SM2a, SM2b). After merging, we normalized the entire dataset to the DHM value corresponding to the first semester of 2016 (which roughly aligns with the summer season in the considered region), to maintain consistency with the artificial thermal stress sequence used in all other results presented in the main paper. However, due to the discrepancies between the historical and projected datasets in the overlapping period, the choice of merging cut-off point (January 2015 vs. January 2025) significantly affects the normalized DHM values for future thermal stress events (cf. Figs. SM2c, SM2d).

We ultimately decided to preserve the full historical dataset up to December 2024 and to append the future projections starting from January 2025. Because the historical dataset exhibits lower thermal stress intensity than the projections in the 2015–2024 interval (Fig. SM2a vs. SM2b), normalizing after a 2025 cut-off leads to higher future DHM values (Fig. SM2c) compared to normalizing after a 2015 cut-off (Fig. SM2d).

This highlights how the merging and normalization strategy—along with the choice of stress proxy (e.g., DHW vs. DHM), climatology baseline, and cumulative stress window—can substantially affect the resulting thermal stress sequence, particularly in terms of intensity values for projected future data. Nonetheless, in our case, it is worth noting that regardless of the merging strategy or the normalization choice, thermal stress values from 2035 onward almost always exceed the critical threshold of 1. This means that, in all cases, the system enters a high-frequency disturbance regime in which corals dominated by thermally sensitive symbionts are pushed beyond their thermal tolerance limits, and thermally tolerant symbionts become dominant across the reef.

This places the system in a dynamical regime analogous to our simulation scenarios with 3-year thermal stress cycles or increasing heatwave intensity. In both cases, the key factor is that the interval between thermal disturbances is shorter than the symbiont reversal time. As a result, the reef remains dominated by thermally tolerant symbionts, allowing it to cope with recurrent stress events, but at the cost of reduced growth. Eventually, the cumulative cost of reduced growth becomes unsustainable, leading to reef extinction.

Finally, we note incidentally that with the chosen merging cut-off point, normalization, climatology threshold and cumulative stress window, our resulting thermal stress sequence successfully recovers the 2016, 2017, and 2020 peaks—also identified in the historical DHW reconstruction by Kamenos and Hennige (2024) for the same geographical area.

### SM2 Simulations without the initial historical sequence

In this section, we present simulations using the same parameters as in the main paper but without the initial historical sequence. The main findings remain consistent with the results shown in the main paper.

#### SM2.1 Increasing intensity scenario

In the increasing intensity scenario, after a transient period of 10 years with no thermal stress, we initiate a sequence of thermal stress events on a 3-year or 5-year cycle, where intensity starts at 0.1 normalized units ($n.u.$) and increases by 0.1 $n.u.$ with each subsequent thermal stress event. Consistent with the results shown in the main paper, we observe that in runs without the initial historical sequence, the counterintuitive effect persists: higher frequencies (3-year cycle) allow for longer reef survival due to the continuous dominance of thermally tolerant symbionts (Fig. SM3, left), compared to lower stress frequencies (5-year cycle) where reversal to thermally sensitive symbionts is favoured, hampering reef survival (Fig. SM3, right).

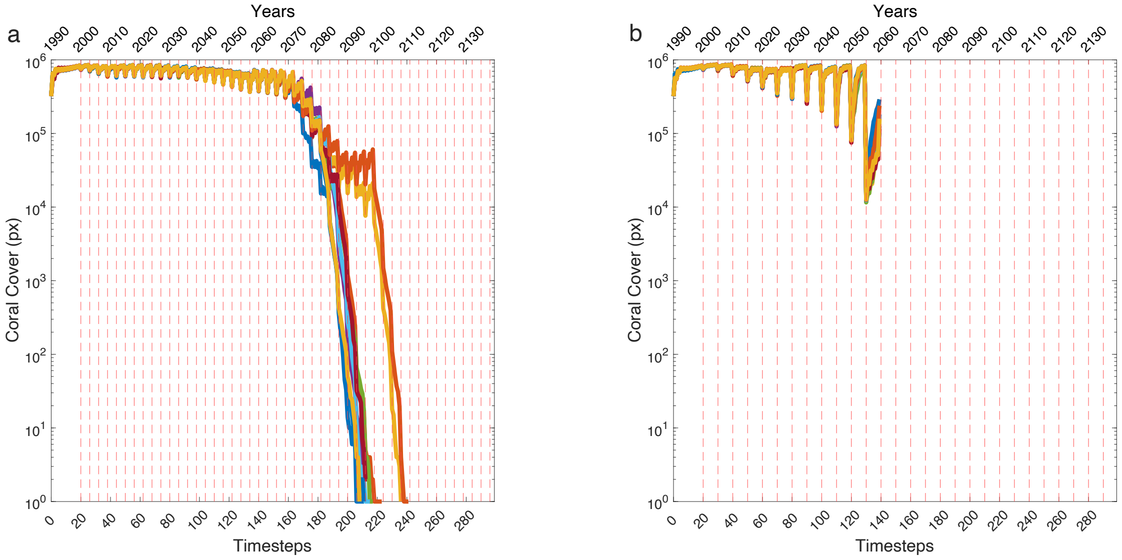


Figure SM3: Projected coral cover over 150 years (300 timesteps) with no initial historical sequence and with increasing intensity thermal stress events every 3 years (left) or every 5 years (right) and abrupt deterministic reversal to sensitive symbionts four years after the last bleaching event. In the 3-year scenario, stress recurs before reversal, maintaining thermal-tolerant dominance and enabling reef persistence since intensities remain sub-lethal for ToS-dominated corals. In the 5-year scenario, reversal completes before the next event, leading to SeS dominance; as stress intensity rapidly becomes lethal for SeS corals (see Table SM5), the reef collapses by 2025. Ten independent runs are shown for each scenario. See Fig.4 in main paper for comparison.

#### SM2.2 Fixed intensity scenario

In the fixed intensity scenario, after a transient period of 10 years with no thermal stress, we initiate a sequence of thermal stress events on a 3-year or 5-year cycle, with intensity kept fixed at 1 $n.u.$ Consistent with the results shown in the main paper, the 3-year thermal stress cycle eventually leads to reef collapse in this scenario due to the reduced growth rate of ToS, which progressively hampers reef structure (Fig. SM3, left). In contrast, the 5-year thermal stress cycle allows for indefinite reef survival in the fixed intensity scenario (Fig. SM3, right), as the improved growth rate of SeS ensures generational turnover of corals. This result is also consistent with the main findings presented in the main paper.

### SM3 Additional reversal dynamics

In all the results presented in the main paper, reversal is deterministic—i.e., it occurs exactly four years after the last thermal stress event (provided no new thermal stress events occur)—and abrupt, meaning that symbiont composition remains unchanged during the four years following

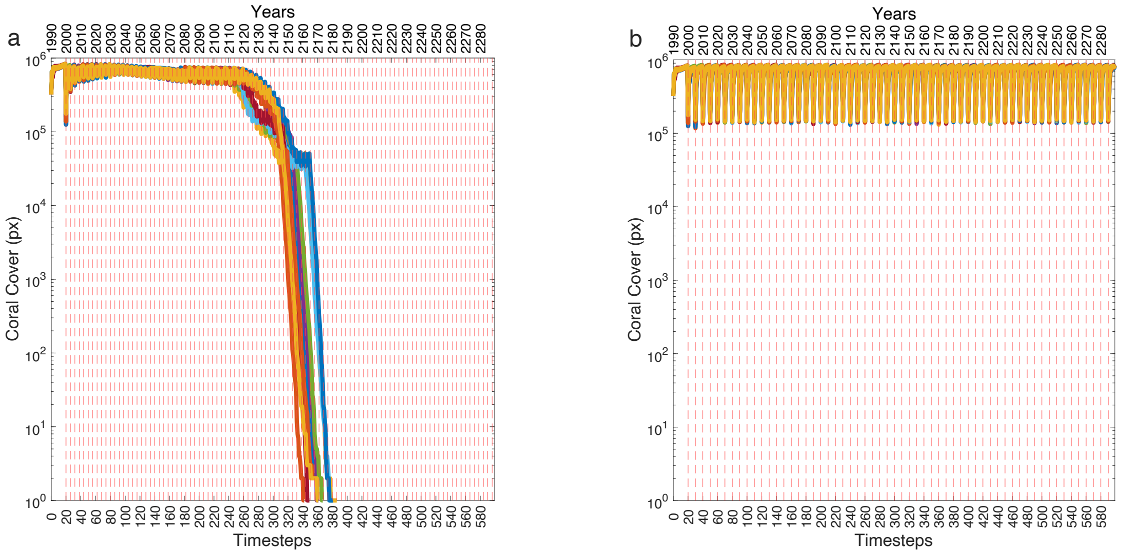


Figure SM4: Projected coral cover over 300 years (600 timesteps) with no initial historical sequence and with fixed intensity thermal stress events every 3 years (left) or every 5 years (right) and abrupt deterministic reversal to sensitive symbionts four years after the last bleaching event. In the 3-year scenario, thermal stress recurs before reversal, maintaining thermal-tolerant dominance, while in the 5-year scenario, reversal completes, resulting in thermal-sensitive dominance. As stress of normalized intensity 1 is sub-lethal for both symbiont types ($P_{\mathrm{mortality}} = 0.1$ for ToS, 0.9 for SeS), both scenarios show initial reef persistence. However, over 250 years (500 timesteps, bottom), only the 5-year scenario supports long-term survival due to the higher growth rates enabled by thermal-sensitive symbionts. Ten independent runs are shown for each scenario. See Fig.5 in main paper for comparison.

the last bleaching event (90% ToS and 10% SeS), reverting back to the pre-stress composition (90% SeS and 10% ToS) afterwards (see Fig. SM7, left).

In this section, we explore two additional reversal dynamics: abrupt probabilistic reversal and gradual deterministic reversal.

#### SM3.1 Abrupt probabilistic reversal

In the main paper, only the switching from thermally sensitive to thermally tolerant symbionts after thermal stress events is modeled probabilistically, while the reversal from thermally tolerant back to thermally sensitive symbionts is deterministic and occurs after a fixed period of four years. Here, we implement fully probabilistic switching, meaning that both the forward switching (from sensitive to tolerant) and the reversal (from tolerant back to sensitive) are stochastic processes.

In this implementation, the reversal time for each coral is drawn from a normal distribution centered at four years with a standard deviation of $\sigma = 0.75$ years, representing variability in how long corals retain thermally tolerant symbionts after the last thermal stress event. Note that reversal remains an abrupt transition—as in the main paper—where symbiont composition shifts instantaneously from 90% thermally tolerant and 10% thermally sensitive symbionts back to 10% tolerant and 90% sensitive once the stress period ends.

In Fig. SM5, we show the results for fully probabilistic switching under an increasing intensity scenario, for both 3-year and 5-year thermal stress cycles. Comparing this figure with Fig. 3 in

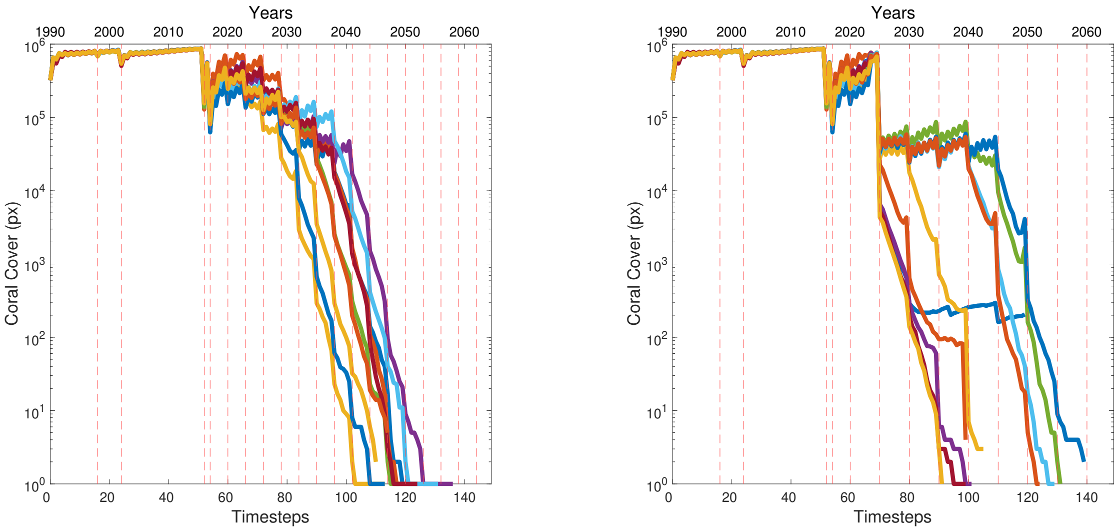


Figure SM5: Projected coral cover over 75 years (150 timesteps) under increasing intensity thermal stress events every 3 years (left) or 5 years (right), with abrupt *probabilistic* reversal to sensitive symbionts occurring on average four years ($\sigma = 0.75$ years) after bleaching. Ten independent runs are shown for each scenario. In the 3-year cycle, results are similar to the abrupt deterministic reversal case (Fig. 4, left), with corals remaining ToS-dominated when the next heatwave arrives, enabling longer reef persistence under increasing stress. In the 5-year cycle, stochastic variability allows some corals to retain ToS dominance when the next heatwave occurs, delaying extinction compared to abrupt deterministic reversal (Fig. 4, right) where all corals revert fully to SeS before the next event, leading to earlier collapse.

the main paper, we observe that under 3-year cycles, results are similar between stochastic and deterministic reversal. However, in the 5-year scenario, probabilistic reversal results in better reef survival than deterministic reversal.

This difference in the 5-year cycle scenario arises because, under deterministic reversal, all corals have already reverted to a thermally sensitive–dominated state when the next thermal stress event occurs after five years, leading to abrupt reef extinction as soon as thermal stress exceeds the tolerance threshold of the sensitive symbiont. In contrast, under probabilistic reversal, some corals are still dominated by thermally tolerant symbionts when the next heatwave arrives, thereby increasing the reef's survival probability due to their higher heat tolerance.

In Fig. SM6, we explore the same probabilistic reversal mechanism under a fixed-intensity scenario. Here, we observe that—compared to the deterministic reversal case, where only the 5-year thermal stress cycle supports reef persistence—stochasticity in reversal timing allows for reef survival even under the 3-year cycle.

This occurs because some corals, due to variability in reversal timing, manage to return to a thermally sensitive–dominated state even within a 3-year cycle. Since thermal stress remains within the survival range of sensitive-dominated corals in the fixed-intensity scenario, these corals can benefit from the enhanced growth associated with thermally sensitive symbionts. Although only a portion of corals effectively revert to sensitive symbiont dominance within three years, in our model and parameterization this proves sufficient to sustain indefinite reef persistence under 3-year thermal stress cycles when abrupt probabilistic reversal is implemented. This contrasts with the 3-year fixed-intensity deterministic reversal scenario, where no corals revert to a sensitive

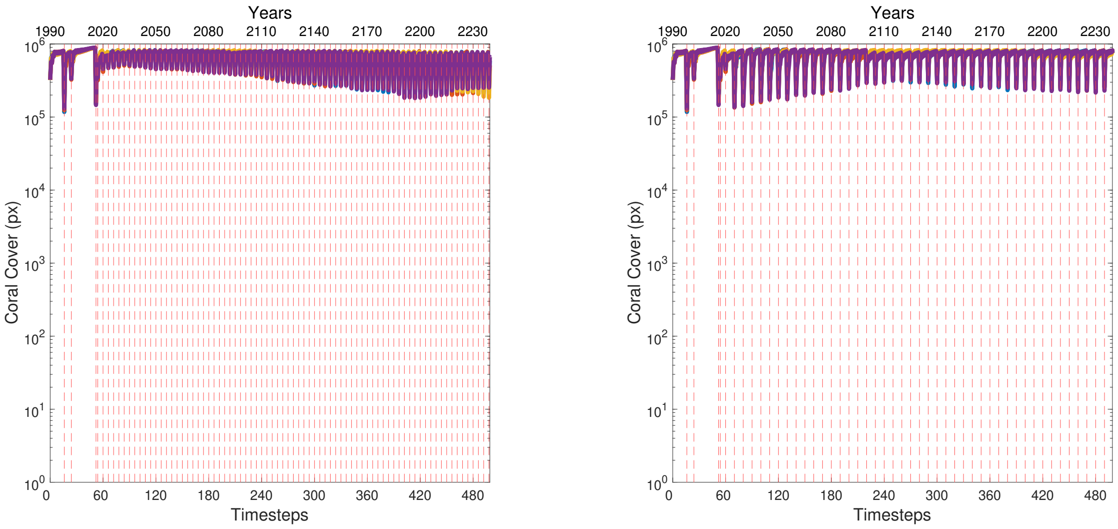


Figure SM6: Projected coral cover over 250 years (500 timesteps) under fixed-intensity thermal stress events every 3 years (left) or 5 years (right), with abrupt *probabilistic* reversal to sensitive symbionts occurring on average four years ($\sigma = 0.75$ years) after bleaching. Four independent runs are shown for each scenario. Compared to the fixed-intensity abrupt *deterministic* reversal case (Fig. 5), where only the 5-year thermal stress cycle supports long-term reef persistence, here probabilistic reversal allows for reef survival even under the 3-year cycle. This occurs because variability in reversal timing enables some corals to revert to SeS dominance within the 3-year cycle, benefiting from higher growth rates while thermal stress remains sub-lethal, thus maintaining reef persistence under conditions that would otherwise lead to collapse in the deterministic scenario.

symbiont state in time, ultimately leading to reef collapse.

#### SM3.2 Gradual deterministic reversal

In simulations with gradual (linear) deterministic reversal, corals gradually revert from thermally tolerant to thermally sensitive symbiont dominance over a characteristic time after bleaching, provided no further thermal stress occurs (Fig. SM7, top and bottom right). We explored scenarios with increasing thermal stress intensity (Figs. SM8, SM10) and fixed thermal stress intensity (Figs. SM9, SM11), under both 3-year and 5-year thermal stress cycles, comparing short (4-year) and long (15-year) gradual reversal periods.

Under increasing intensity, a short 4-year gradual reversal (Fig. SM8) leads to a rapid increase in thermally sensitive symbionts within corals, making them vulnerable when thermal stress intensities exceed the tolerance threshold for sensitive-dominated corals. Consequently, reefs fail to survive the 2017 event under both the 3-year and 5-year cycles. In contrast, a slower 15-year gradual reversal (Fig. SM10) results in a shallower reversal slope, allowing corals to retain a substantial proportion of thermally tolerant symbionts when new heatwaves occur. This enables reefs to survive the 2017 event and maintain persistence for a longer period. Reef persistence occurs even under the 5-year cycle, in contrast to the abrupt 4-year reversal scenario (see Fig. 4, right panel, in the main paper), where reefs collapse almost immediately due to full sensitive-symbiont dominance.

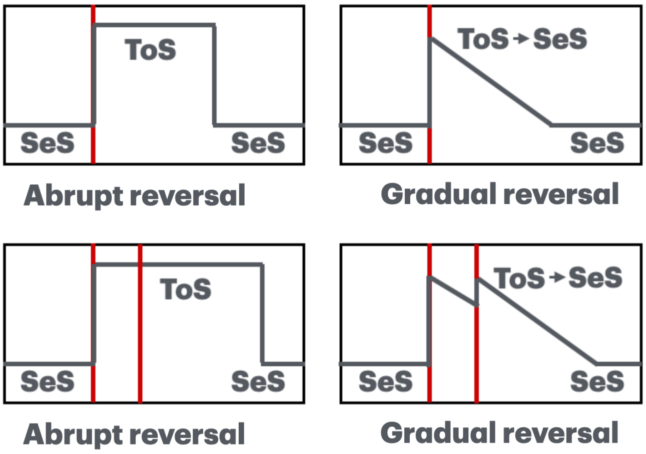


Figure SM7: Top left: in the 4-year abrupt reversal scenario, when a thermal stress occurs symbiont composition becomes ToS-dominated for 4 years before abruptly reverting to SeS dominance if no additional thermal stress occurs. Bottom left: in the abrupt reversal scenario, if a new thermal stress event occurs during ToS dominance, then this is extended for another 4 years. Top right: in the gradual reversal scenario, corals switch to ToS dominance after a thermal stress event, then the ToS→SeS immediately starts and takes place over a characteristic time, provided no additional thermal stresss events occur. Bottom right: in the gradual reversal scenario, if a new thermal stress event occur, then maximum allowed ToS dominance (90% in our model) is recovered, then the ToS→SeS reversal process starts again.

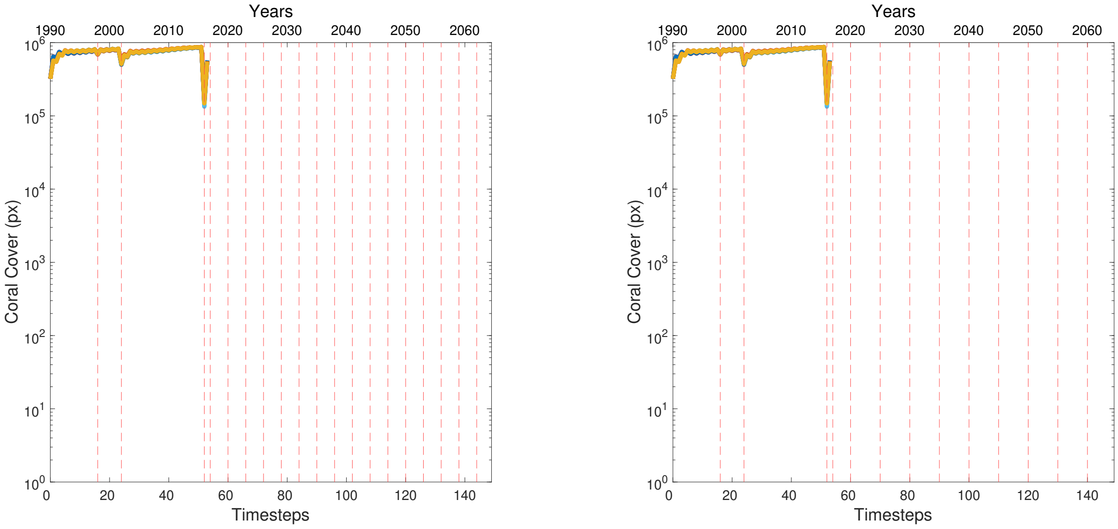


Figure SM8: Projected coral cover over 75 years (150 timesteps) under increasing intensity thermal stress events every 3 years (left) or 5 years (right), with *gradual* (linear) deterministic reversal to sensitive symbionts occurring over four years after bleaching. With gradual reversal, the steep reversal slope rapidly increases the SeS proportion within corals, leading to early collapse when thermal stress intensities become lethal for SeS-dominated corals. Consequently, under these settings, reefs fail to survive the 2017 event in both the 3-year and 5-year thermal stress cycles. Ten independent runs are shown for each scenario.

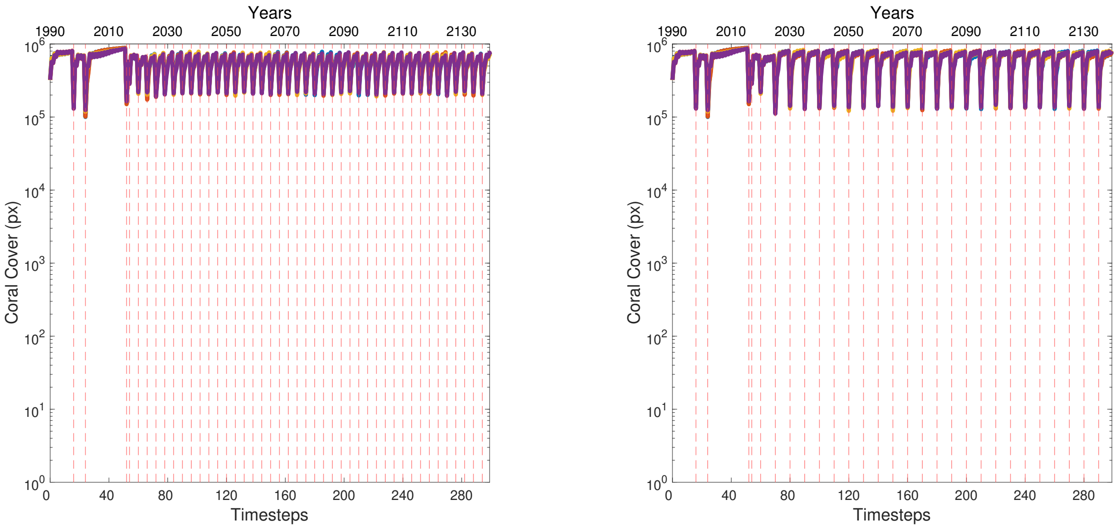


Figure SM9: Projected coral cover over 150 years (300 timesteps) under fixed intensity thermal stress events every 3 years (left) or 5 years (right), with *gradual* (linear) deterministic reversal to sensitive symbionts occurring over four years after bleaching. Gradual reversal allows for hybrid SeS/ToS coral dominance thoughout the reef. As we are in the fixed intensity scenario both SeS and ToS-dominated corals are below thermal tolerance threshold, so that the reef can exploit both SeS-endowed and ToS-endowed benefits (increased growth and increased thermal resilience respectively). Four independent runs are shown for each scenario.

Under fixed intensity, gradual reversal allows the reef to maintain a hybrid symbiont composition throughout the cycle (Figs. SM9, SM11). Since thermal stress intensities remain below the tolerance thresholds for both sensitive- and tolerant-dominated corals, the reef can simultaneously benefit from the increased growth rates conferred by sensitive symbionts and the thermal resilience provided by tolerant symbionts. This hybrid composition enhances reef stability and persistence under repeated thermal stress events in both the 3- and 5-year cycles.

### SM4 Additional Supporting Results

In this section, we present additional simulations exploring how changes in key parameters—specifically the symbiont switching probability ($P_{\text{switching}}$) and symbiont-endowed growth rates—affect coral reef persistence under fixed-intensity thermal stress events with 3-year and 5-year cycles.

#### SM4.1 Effect of varying symbiont switching probability.

In the standard configuration used in the main paper, $P_{\text{switching}} = 0.8$ ensures that the majority of corals switch from thermally sensitive to thermally tolerant symbionts after each thermal stress event, with deterministic reversal to a thermally sensitive-dominated state after four years. As at fixed intensity we are within thermal tolerance levels for both symbionts, this configuration supports indefinite reef persistence in the 5-year cycle scenario. Thermal resilience offered by thermally tolerant symbionts is never exploited, because of deterministic reversal to sensitive symbionts before the next thermal stress event. However, the robust growth provided by full occupation of thermally sensitive symbionts in the last year of the cycle (plus partial sensitive

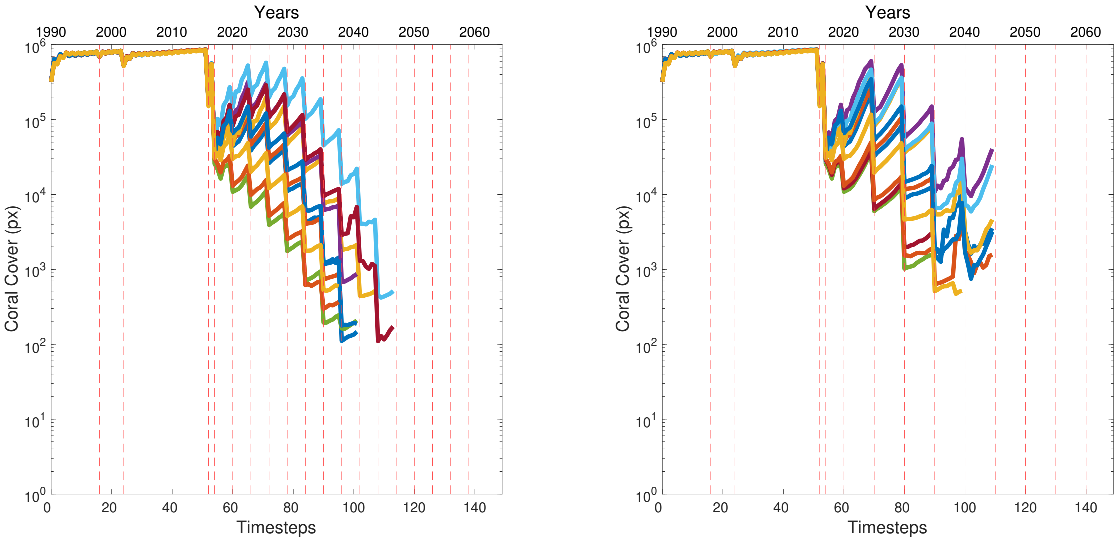


Figure SM10: Projected coral cover over 75 years (150 timesteps) under increasing intensity thermal stress events every 3 years (left) or 5 years (right), with gradual linear reversal to sensitive symbionts occurring over fifteen years after bleaching. When gradual reversal occurs at a slower pace, a significant proportion of corals are still ToS-dominated when the new heatwave occurs. This enables corals to survive the 2017 event and maintain reef persistence for longer, even under the 5-year thermal stress cycle, compared to the abrupt 4-year reversal case where reefs collapse earlier under the same conditions (Fig. 4, right panel, in the main paper). Ten independent runs are shown for each scenario.

symbiont persistence even in the first four years due to the probabilistic nature of switching) allows the reef to survive indefinitely (Fig. 5, bottom right and Fig. 6d in the main paper). The presence of thermally tolerant symbionts in this configuration serves the purpose of protecting the reef in case of thermal perturbations at higher intensity in the first four years of each cycle, enhancing reef's robustness. In contrast, in the 3-year cycle scenario (Fig. 5, bottom left, and Fig. 6c) the reef becomes soon dominated by thermally tolerant symbionts only. This increases survival at each thermal stress events, but permanently reduces growth to an extent that causes reef extinction. In that regard full thermally tolerant dominance can be seen in our model as a temporary emergency response that cannot sustain alone reef survival in the long term.

Disabling symbiont switching ($P_{\text{switching}} = 0$) entirely results in full, continuous dominance by thermally sensitive symbionts. This enhances coral growth so that the reef can survive indefinitely in both the 3-year and 5-year cycle (see Fig. SM12). However exclusive sensitive symbiont dominance leaves the reef highly vulnerable to thermal disturbances, with total collapse occurring as soon as stress intensities exceed the tolerance limits of sensitive symbionts.

At the opposite extreme, forcing $P_{\text{switching}} = 1$ leads to complete switching to thermally tolerant symbionts after each stress event. Therefore in the 5-year cycle scenario reversal to thermally sensitive symbionts only occurs in the final year before the next event. This configuration maintains reef survival (Fig. SM13, although with lower overall coral cover compared to the $P_{\text{switching}} = 0.8$ scenario shown in the main paper, where sensitive symbiont partially remain also in the first four years. This demonstrates that in our model even a short period of thermal sensitive dominance in the last year of the cycle can sustain reef persistence, and that sensitive symbiont persistence in the first four years also plays a secondary but noticeable role

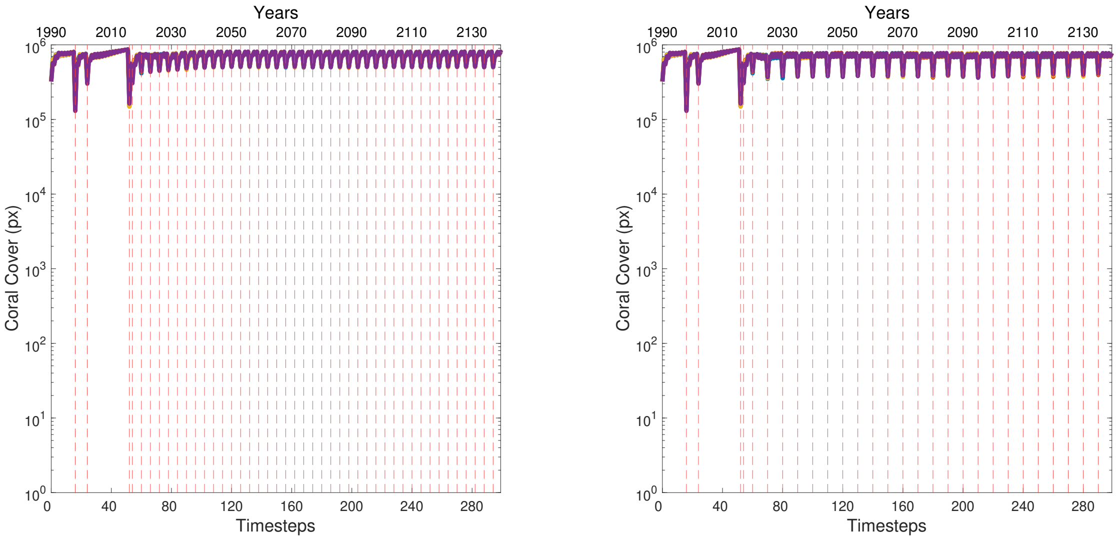


Figure SM11: Projected coral cover over 150 years (300 timesteps) under fixed intensity thermal stress events every 3 years (left) or 5 years (right), with gradual linear reversal to sensitive symbionts occurring over fifteen years after bleaching. Gradual reversal allows for hybrid SeS/ToS coral dominance thoughout the reef. As we are in the fixed intensity scenario both SeS and ToS-dominated corals are below thermal tolerance threshold, so that the reef can exploit both SeS-endowed and ToS-endowed benefits (increased growth and increased thermal resilience respectively). Note that the only difference with the scenario with gradual reversal over four years is that here less coral cover is lost after thermal stress, because there are more ToS-dominated corals due to a slower reversal process. Four independent runs are shown for each scenario.

in enhancing reef's stability. In contrast in the 3-year cycle scenario the reef can never benefit from sensitive symbiont-driven robust growth, leading to reef extinction.

When corals are forced to remain dominated by thermally tolerant symbionts continuously, without any reversal, the reef ultimately collapses in both the 3- and 5-year cycle (Fig. SM14). This highlights the critical role that even brief periods of dominance by thermally sensitive symbionts play in out model in maintaining coral growth and long-term reef stability under recurring stress.

#### SM4.2 Effect of symbiont-endowed growth rates.

To further test the role of growth trade-offs, we conducted simulations where SeS and ToS endow identical growth rates. When SeS growth is reduced to match the low ToS growth rate of 0.1 (Fig. SM15), the reef collapses rapidly due to the absence of SeS-driven enhanced growth, demonstrating the importance of growth advantages provided by SeS. Conversely, when ToS growth is increased to match the high SeS rate of 0.9 (Fig. SM16), the reef maintains high resilience while preserving thermal stress robustness, as ToS persistence no longer hampers growth. Although this scenario represents an unrealistic removal of the growth trade-off, it highlights that the optimal configuration in our model remains the default scenario (Fig. 7d in the main paper), where ToS provides resilience against frequency perturbations while SeS ensures efficient coral growth.

Together, these additional supporting results illustrate how the interplay between symbiont switching probabilities, symbiont-endowed growth rates, and deterministic reversal critically shapes reef persistence and resilience under recurring thermal stress, providing mechanistic insight into the bet-hedging strategies observed in our model under different parameter regimes.

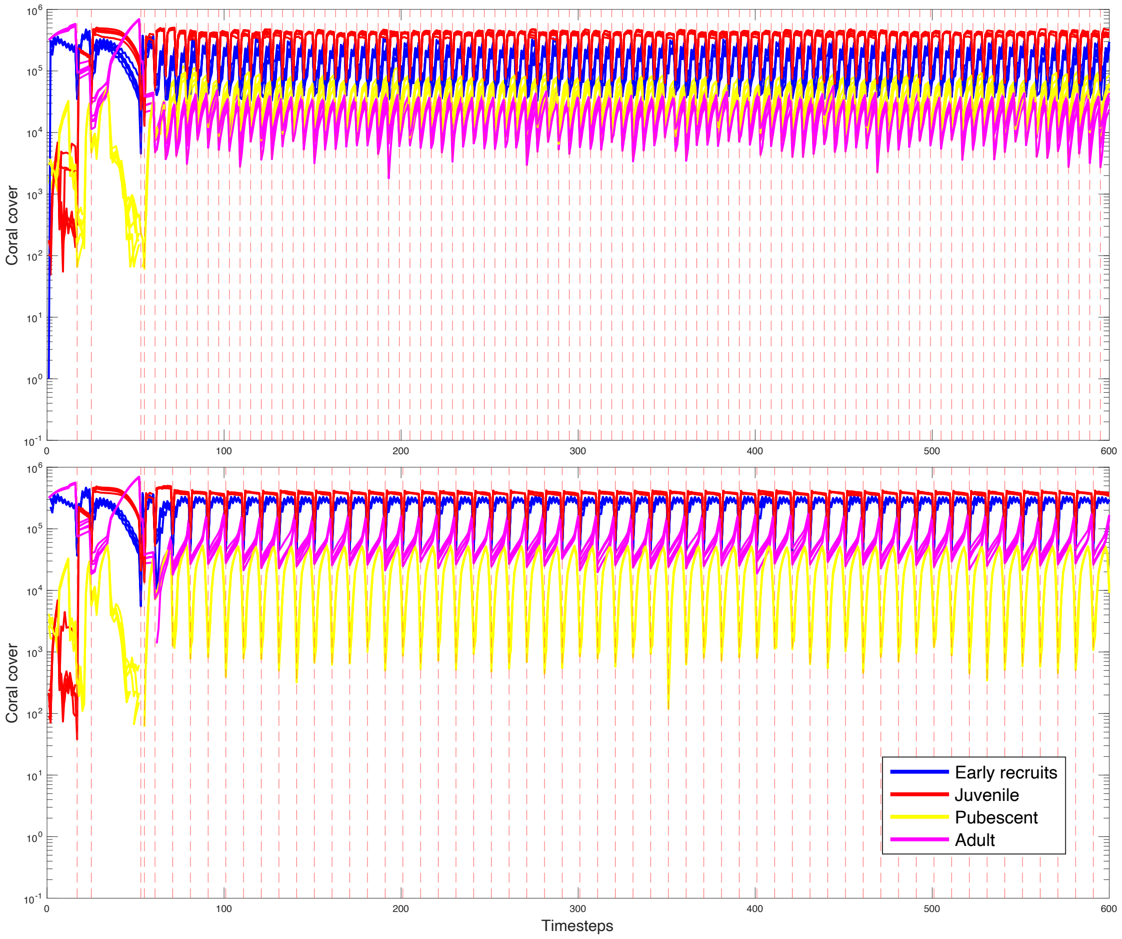


Figure SM12: Projected coral cover over 300 years (600 timesteps) for different life stages under fixed-intensity thermal stress events occurring every 3 years (top) or 5 years (bottom), with $P_{switching} = 0$. Corals do not switch from thermally sensitive to thermally tolerant symbionts after stress events, remaining fully dominated by thermally sensitive symbionts throughout. This removes the growth penalty associated with thermally tolerant symbiont dominance and allows for additional growth in both the 3- and 5-year cycles, enhancing reef persistence. However, compared to the scenario with $P_{switching} = 0.8$, these reefs are highly unstable under thermal disturbances: due to full sensitive symbiont dominance, the entire reef becomes extinct at the first thermal stress event exceeding the symbionts' tolerance threshold ($T > 1$ $n.u.$). Five independent runs are shown.

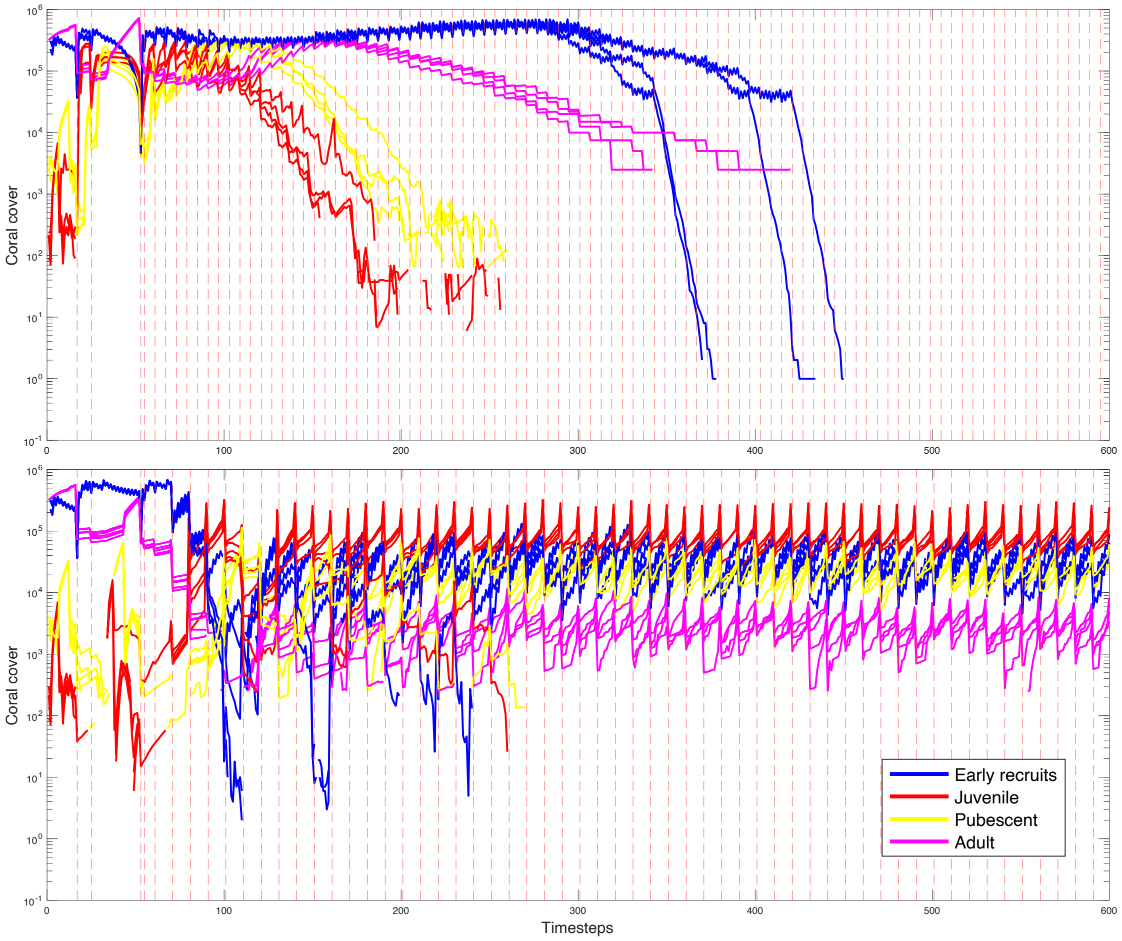


Figure SM13: Projected coral cover over 300 years (600 timesteps) for different life stages under fixed-intensity thermal stress events occurring every 3 years (top) or 5 years (bottom), with $P_{switching} = 1$. All corals switch from thermally sensitive to thermally tolerant symbionts after stress events. This means that in the 5-year cycle, corals are fully dominated by thermally tolerant symbionts for the first four years of each cycle and revert to thermally sensitive dominance in the final year, while in the 3-year cycle, corals remain fully dominated by thermally tolerant symbionts throughout. In the 5-year cycle, reversal to thermally sensitive dominance in the last year allows for enhanced growth, protecting the reef from extinction. In contrast, in the 3-year cycle, continuous dominance by thermally tolerant symbionts eventually leads to reef extinction. Four and nine independent runs are shown, respectively.

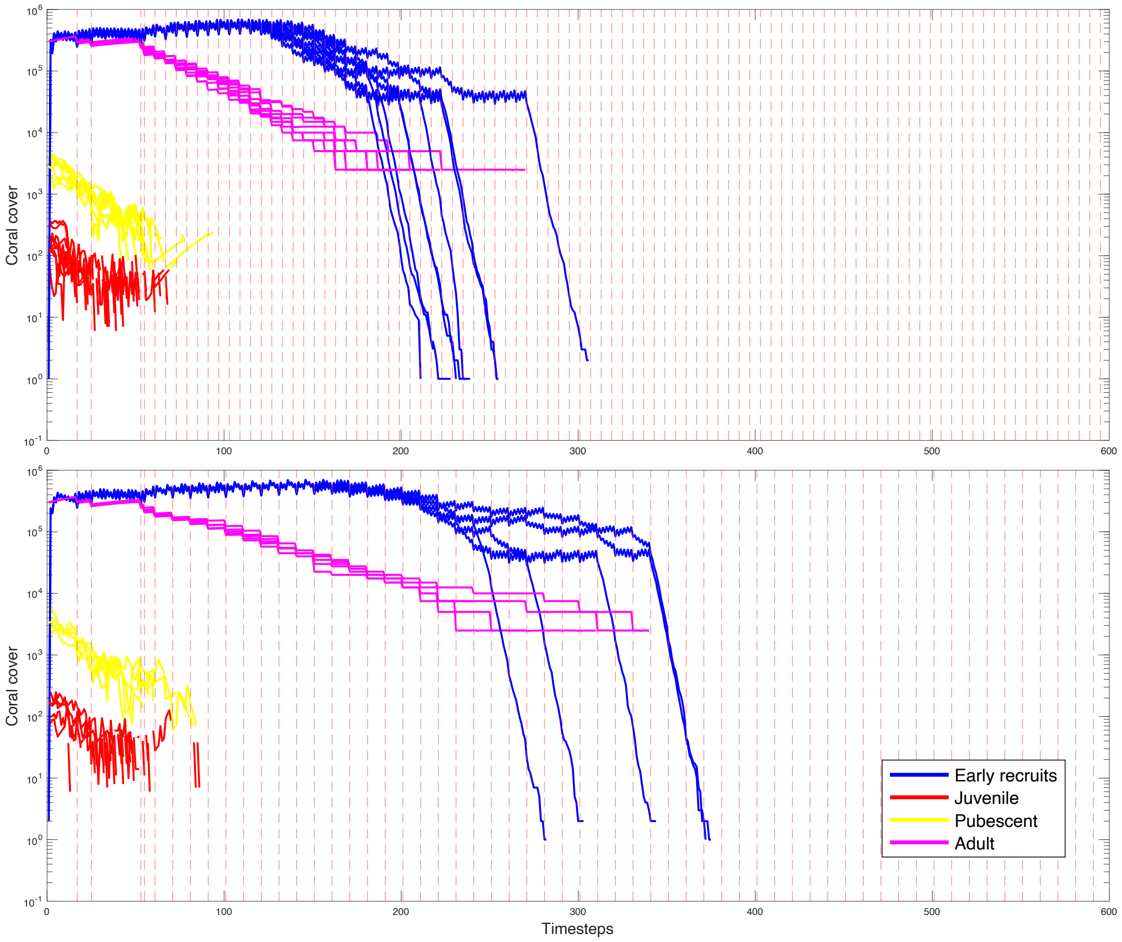


Figure SM14: Projected coral cover over 300 years (600 timesteps) for different life stages under fixed-intensity thermal stress events occurring every 3 years (top) or 5 years (bottom), with exclusive dominance by thermally tolerant symbionts. Although recruit production initially masks reef collapse, continuous dominance by thermally tolerant symbionts eventually leads to reef extinction in both the 3- and 5-year scenarios. Since the only difference between this set of runs and the setup with $P_{switching} = 1$ is the presence of thermally sensitive symbiont dominance during the last year of the 5-year cycle, this confirms that enhanced coral growth endowed by thermally sensitive symbionts in that final year enables reef survival when $P_{switching} = 1$. Nine and five independent runs are shown, respectively.

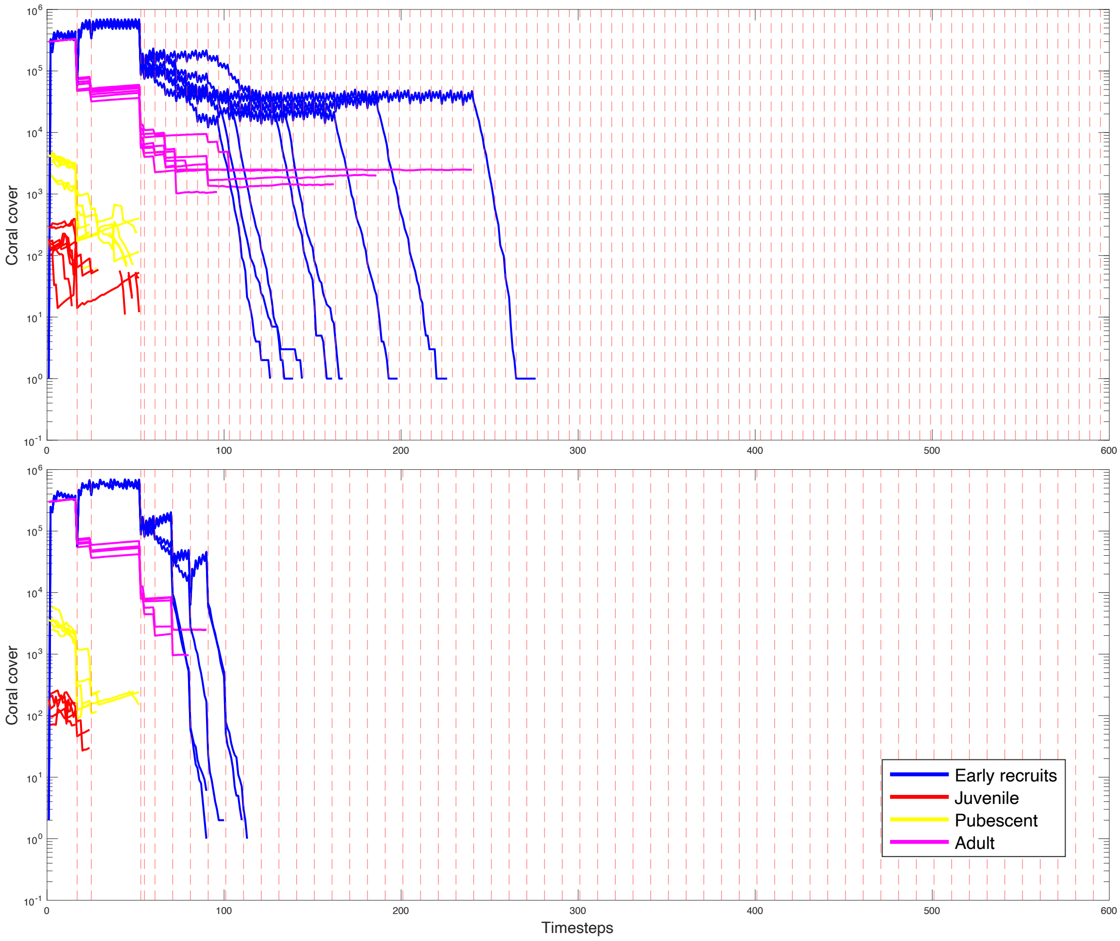


Figure SM15: Projected coral cover over 300 years (600 timesteps) for different life stages under fixed-intensity thermal stress events occurring every 3 years (top) or 5 years (bottom), with $P_{switching} = 0.8$ and the growth rate endowed by thermally sensitive symbionts reduced to 0.1 (as low as that for thermally tolerant symbionts). This represents one of the worst possible scenarios, as there is no enhanced growth benefit from thermally sensitive symbionts, and all corals revert to sensitive symbionts before the stress event in the 5-year cycle scenario, leaving the reef unprotected, which leads to early extinction. Continuous dominance by thermally tolerant symbionts in the 3-year scenario provides some thermal protection and extends reef survival, but the reef ultimately collapses. Five independent runs are shown.

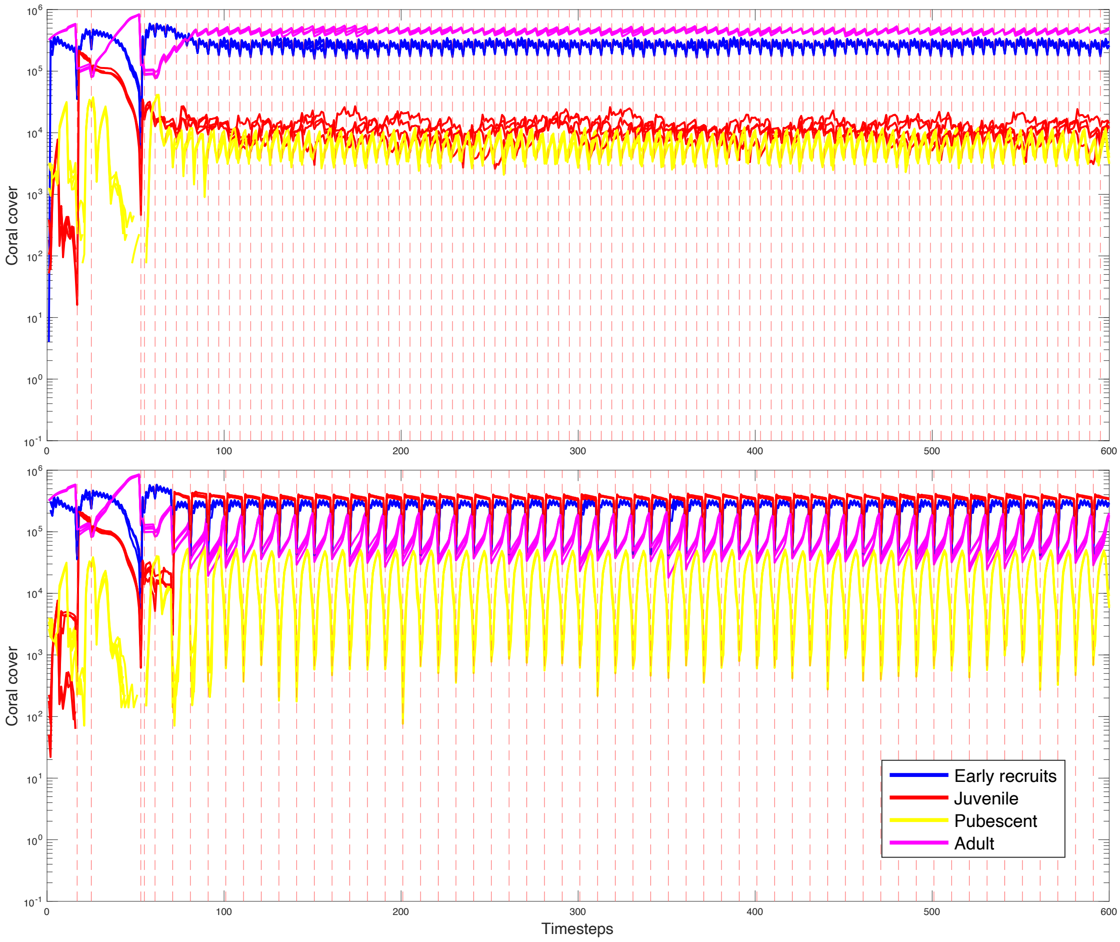


Figure SM16: Projected coral cover over 300 years (600 timesteps) for different life stages under fixed-intensity thermal stress events occurring every 3 years (top) or 5 years (bottom), with $P_{switching} = 0.8$ and the growth rate endowed by thermally tolerant symbionts increased to 0.9 (as high as that for thermally sensitive symbionts). This represents one of the best possible scenarios, as switching to thermally tolerant symbionts does not hamper coral growth. In the 5-year scenario, the reef is dominated by thermally sensitive symbionts when the next thermal stress event occurs, leading to partial reef mortality, which is subsequently recovered due to high growth levels. In the 3-year scenario, mortality is negligible due to thermal tolerant symbiont dominance, and high growth levels allow for long-term reef persistence. Notably, this is the only case in our model in which adult corals consistently dominate over other life stages throughout the simulation.

Table SM4: Summary of general model parameters across different versions of the reef model.

| **Parameter** | **Additional description** | Mumby et al. (2007) | Ortiz et al. (2013) | **Current model** |
|---|---|---|---|---|
| Number of coral types | - | 2 (1 'brooder' type and 1 'spawner' type). Brooders and Spawners only differ in terms of growth, and of larval production during recruitment. | as in Mumby 2007? | 1 (generic type) |
| Max number of corals per cell | - | 3 (in total, between brooders and recruiters, all stages) | - | unlimited (i.e., max 2500 individuals per cell, constrained only by turf presence) |
| Hurricane mortality | - | Yes | No | No |
| Thermal stress mortality | - | No | Yes | Yes |
| Reef size | - | 2500 cells of $0.25m^2$ each | 400 cells of $0.25m^2$ each | 400 cells of $0.25m^2$ each |
| Time interval | - | 6 months | 6 months | 6 months |
| Symbionts incorporation | Inclusion of symbiont type in the model | No | Yes | Yes |
| Initial symbiont composition in the reef | % of corals in the reef that are dominated by a given symbiont type | - | 5% ToS, 95% SeS | 100% SeS (we assume No recent heatwaves, and that after a given time without significant thermal stress events all corals are dominated by SeS) |
| Thermal-induced symbiont switching | Sensitive → Tolerant | No | Yes | Yes |
| Thermal-induced symbiont reversal | Tolerant → Sensitive | No | No | Yes |
| Linear interpolation between symbiont properties | Whether the symbiont-endowed characteristic of the coral depend only on dominant symbiont type or on the relative composition of symbiont community | - | No | Yes |

*Continued on next page*

Table SM4 – *Continued from previous page*

| **Parameter** | **Additional description** | Mumby et al. (2007) | Ortiz et al. (2013) | **Current model** |
|---|---|---|---|---|
| Corals growth | Lateral extension rate of cross-sectional basal area | Brooders: $G_r^b = 0.8$ cm yr$^{-1}$ Spawners: $G_r^b = 0.5$ cm yr$^{-1}$ | as in Mumby 2007? | Basal growth rate $G_r^b$ is the lateral extension rate of cross-sectional basal area; $G_r^b$ depends on symbiont composition; Tolerant symbiont $\rightarrow G_r^b = 0.1$; Sensitive symbiont $\rightarrow G_r^b = 0.9$; $G_r^b$ can be reduced due to coral-MA competition |
| Direct coral-MA competition for juvenile corals (area $< 60cm^2$) | Growth rate $G_R$ is reduced to $G_R^*$ in presence of macroalgae | if $M_{4cells} > 80\%$ then $G_r^e = 0 \times G_r^b$ ; if $60\% < M_{4cells} \leq 80\%$ then $G_r^e = 0.3 \times G_r^b$ | as in Mumby 2007? | Same as in Mumby 2007 |
| Direct coral-MA competition for pubescent and adult corals (area $\geq 60cm^2$) | Growth rate $G_R$ is reduced to $G_R^*$ in presence of macroalgae | if $M_{4cells} \geq 60\%$ then $G_r^e = 0.5 \times G_r^b$ | as in Mumby 2007? | Same as in Mumby 2007 |
| Corals competitive exclusion | Growth of corals in absence of available space (turf) | ? | ? | If there is not enough turf for all corals to grow, then the biggest coral grows at the expense of the smaller ones. |
| Vegetative macroalgal growth ($MA \rightarrow CA$) | Vegetative growth of MA over cropped algae (CA) | If $C_{4cells} < 0.5$ then $P_{CA \rightarrow MA} = M_{4cells}$ ; if $C_{4cells} \geq 0.5$ then $P_{CA \rightarrow MA} = 0.75 \times M_{4cells}$ , where $C_{4cells}$ is the % of coral in the 4-VN neighborhood (i.e., MA - Coral competition reduces vegetative MA overgrowth on CA) | as in Mumby 2007? | Same as in Mumby 2007 |
| Grazing by parrotfish | Maximum % of turf grazed by parrotfish in one semester | 40% $\times$ 6 months; no distinction between algal states (CA & MA); parrotfish and urchin can graze same areas | as in Mumby 2007? | 80% $\times$ 6 months |

*Continued on next page*

Table SM4 – *Continued from previous page*

| **Parameter** | **Additional description** | Mumby et al. (2007) | Ortiz et al. (2013) | **Current model** |
|---|---|---|---|---|
| Grazing by sea urchin | Maximum % of turf grazed by sea urchin in one semester | 53% $\times$ 6 months; no distinction between algal states (CA & MA); parrotfish and urchin can graze same areas | as in Mumby 2007? | - |
| Larval predation by parrotfish | Coral mortality due to PF | 15% recruits (area $\leq$ 5 cm$^2$ every 6 months) | as in Mumby 2007? | Same as in Mumby 2007 |
| Algal aging | - | $P_{CA \rightarrow MA}$ = 70% after 6 months from grazing; $P_{CA \rightarrow MA}$ = 100% after 12 months from grazing | as in Mumby 2007? | $P_{CA \rightarrow MA} = 5\%$ after 6 months from grazing; $P_{CA \rightarrow MA}$ = 100% after 12 months from grazing |
| Coral reproduction | - | Excluded, assume constant rate of coral recruitment from outside reef (i.e. no stock-recruitment dynamics). Recruitment rate is 2 recruits per $0.25m^2$ per semester (brooders), 0.2 recruits per $0.25m^2$ per semester (spawners). Adjusted for rugosity. | ? | Reproduction occurs every 12 months. Stock-recruitment dynamics. # of larvae = $F_C^e \times A_C$ where $A_C$ is the cross-sectional basal area of coral and effective fecundity of coral $F_C^e$ depends on MA % in $M_{4cells}$: $F_C^e = 0.75 \times F_C$ if $M_{4cells} > 0.5$. Corals can produce more larvae than available space on reef. |
| Whole colony mortality | Random whole colony mortality | 2% every 6 months for pubescent corals $(60-250cm^2)$; 1% every 6 months for adult corals $(\geq 250cm^2)$ | ? | 2% every 6 months for pubescent corals $(60-250cm^2)$ not implemented for adults |
| Partial colony mortality | Random partial colony mortality | Ppm $= 1 - [60 + (-12\ln(\chi))]$; $\ln[(\text{Apm} \times 100) + 1] = -0.5 + (1.1\ln(\chi))$; where Ppm is the probability of a partial mortality event, Apm is the area of tissue lost in a single event, and $\chi$ is the size of the coral in $cm^2$. (unclear) | ? | No |

*Continued on next page*

Table SM4 – *Continued from previous page*

| **Parameter** | **Additional description** | Mumby et al. (2007) | Ortiz et al. (2013) | **Current model** |
|---|---|---|---|---|
| Macroalgal Overgrowth ($MA \rightarrow Coral$) | MA overgrowth over corals | $O_{C\rightarrow M} = M_{4cells} \cdot P_i \cdot 4/7$ where $M_{4cells}$ is the % of MA in the VM 4-cell neighbourhood, $P_i$ is the perimeter of the cross-sectional basal area of coral, 4/7 is an experimental scaling factor | as in Mumby 2007? | Same in Mumby 2007 |
| Direct coral-MA competition for pubescent and adult corals (area $\leq 60cm^2$) | - | if $M_{4cells} > 60\%$ then $G_r^e = 0 \times G_r^b$ | as in Mumby 2007? | Same as in Mumby 2007 |
| $P_{\text{mortality}}$ due to thermal stress | - | - | $P_{\text{mort}.ToS} = 0.7 \times P_{\text{mort}.SeS}$; $P_{\text{mort}.SeS} =?$ | $P_{\text{mortality}ToS} = 10\%$ when 100% ToS-dominated; $P_{\text{mortality}SeS} = 90\%$ when 100% SeS-dominated (for mixed symbiont composition values are interpolated) |
| $P_{\text{switching}}$ due to thermal stress | $P(\text{SeS} \rightarrow \text{Tosswitching})$ | - | 40% | 80% |

*Continued on next page*

| **Parameter** | **Additional description** | Mumby et al. (2007) | Ortiz et al. (2013) | **Current model** |
|---|---|---|---|---|
| Reversal | - | - | - | Abrupt deterministic scenario: ToS $\rightarrow$ SeS reversal occurs with P=1 exactly after 8 semesters from the last thermal stress event. If an additional thermal stress event occurs in the meanwhile, ToS permanence is extended for an additional 4 years. Symbiont composition is fixed at 90% Tos, 10% Ses after thermal stress, or at 10% Tos, 90% Ses during thermally stable periods (see Fig.SM7). Gradual deterministic scenario: Same as abrupt deterministic scenario, except for the fact that reversal towards sensitive symbiont is gradual and begins immediately after the onset of ToS dominance due to thermal stress events. Abrupt stochastic scenario: Same as abrupt deterministic scenario, except for the fact that ToS $\rightarrow$ SeS occurs with a different characteristic time for each coral, normally distributed around 4 $\pm\sigma = 0.75$ years. |

Table SM4: Summary of general model parameters across different versions of the reef model. The table compares key structural and ecological assumptions from Mumby et al. (2007), Ortiz et al. (2013), and the current model implementation. Comparison with Mumby (2006) is omitted due to space constraints.
